# Secure LSL: A Unified Encryption Architecture for the Lab Streaming Layer

**DOI:** 10.64898/2026.07.07.737068

**Authors:** Seyed Yahya Shirazi, Scott Makeig

**Affiliations:** Swartz Center for Computational Neuroscience, Institute for Neural Computation, University of California San Diego, La Jolla, CA, United States

**Keywords:** biosignal security, neural recording, mobile brain/body imaging (MoBI), brain behavior quantification and synchronization (BBQS), regulatory compliance, real-time data streaming

## Abstract

**Objective:** The Lab Streaming Layer (LSL) protocol, widely adopted for synchronized multimodal biosignal recording in neuroscience research, transmits all data in plaintext, exposing sensitive neural and physiological recordings to interception and tampering and creating regulatory liability for clinical and commercial deployments across most major international jurisdictions. We present Secure LSL, the unified encryption architecture for the protocol: a novel security layer that authenticates devices and encrypts biosignal streams through transparent, drop-in modifications to the core library, requiring no application changes and no recompilation for dynamically linked clients.

**Approach:** We implement encryption at the *liblsl* core library level using a shared keypair authorization model with ChaCha20-Poly1305 authenticated encryption. All authorized devices share a common Ed25519 keypair, and public key verification during connection establishment ensures only authorized devices communicate. The architecture enforces network-wide security consensus, requiring all connected devices to operate in either secure or insecure mode, eliminating vulnerable mixed environments, and operates transparently with zero code changes to existing applications.

**Main Results:** The architecture preserves application programming interface (API) transparency, so existing applications need no code changes (legacy devices must update to connect to secured outlets). Across five hardware platforms spanning x86 desktop, Apple Silicon laptop, embedded ARM single-board, and Xtensa microcontroller targets, encryption adds sub-millisecond latency in all desktop and embedded ARM configurations, with overhead in the single-digit percent range (approximately 4 to 9%, the lowest values within measurement noise of zero) for typical 64-channel, 1000-Hz configurations. A clean-room ESP32 implementation extends transparent encryption to dual-core microcontrollers with no measurable push-path overhead and approximately 2 kB additional static random-access memory (SRAM) consumption, enabling secured wearable and ambulatory biosensor deployments.

**Significance:** By implementing security within the protocol core rather than requiring application-level changes, we transform LSL from a research-only protocol to a security-capable platform for clinical settings, multi-institution collaborations, and commercial products, while preserving its zero-configuration philosophy.

## 1 Introduction

### 1.1 Lab Streaming Layer and Its Security Gap

The Lab Streaming Layer (LSL) has become the *de facto* standard for synchronized multimodal data acquisition in neuroscience research, enabling millisecond-precision synchronization across heterogeneous biosignal sources without a common clocking mechanism, including electroencephalography (EEG), eye tracking, motion capture, and other physiological sensor systems [1]. LSL’s publisher-subscriber architecture uses User Datagram Protocol (UDP) multicast for service discovery and Transmission Control Protocol (TCP) for data transmission, prioritizing ease of use through zero-configuration networking. This design philosophy has proven successful, with LSL now supporting over 150 hardware driver applications and establishing interoperability across client software written in C/C++, Python, MATLAB, Java, C#, and several other programming languages.

LSL was designed for trusted laboratory-scale local networks where security was not a requirement. Its official documentation acknowledges that “the SessionID is not a security feature… you are still able to intercept packets involved in a session that is not yours” [2]: all biosignal data transmits in plaintext over TCP, UDP multicast broadcasts expose stream metadata to entire networks, and no authentication prevents unauthorized access. Yet LSL’s adoption has expanded well beyond this original scope, with commercial neurotechnology products, mobile EEG devices communicating with phones or base stations, industrial systems, and clinical brain-computer interfaces now relying on it, often across networks with external connectivity. None of the ecosystem’s applications provide security features, as the platform was not designed for them.

### 1.2 Regulatory Implications

Most LSL-compatible devices are research-grade instruments (EEG amplifiers, eye trackers, motion capture systems) rather than Class II/III medical devices, and are therefore exempt from medical-device cybersecurity requirements such as the European Union Medical Device Regulation (MDR) or United States Food and Drug Administration (FDA) 510(k) requirements. Region-specific data-protection regulations, however, apply regardless of device classification. Across major international jurisdictions, including the Health Insurance Portability and Accountability Act (HIPAA) in the United States [3]; the General Data Protection Regulation (GDPR) [4], the Network and Information Security Directive 2 (NIS2) [5, 6], the European Health Data Space (EHDS) [7], and the Cyber Resilience Act (CRA) [8] in the European Union; the Personal Information Protection Law (PIPL) and Cybersecurity Law (CSL) in China [9, 10]; the Act on the Protection of Personal Information (APPI) in Japan [11]; and the Personal Information Protection Act (PIPA) in South Korea [12], transport-layer encryption of identifiable biosignal data is either explicitly required or treated as expected practice. Plaintext LSL transport is therefore incompatible with clinical environments, multi-institution collaborations involving identifiable participant data, and commercial neurotechnology products serving these jurisdictions. Neural and biosignal data are, moreover, increasingly recognized as a distinct category warranting heightened protection: a growing “neurorights” scholarship argues that general data-protection frameworks, including the GDPR, may not by themselves be sufficient for neurotechnology [13, 14, 15], which strengthens rather than weakens the case for encrypting these streams by default. Section 5.6 details these obligations and how Secure LSL addresses them.

### 1.3 Design Constraints and Requirements

Securing LSL presents challenges arising from the protocol’s architecture and ecosystem structure. The ecosystem comprises approximately 150 hardware-specific applications coded in several programming languages, most driving specific data-source instruments (EEG amplifiers, eye trackers, motion capture systems) that operate autonomously or in embedded contexts. Updating these diverse applications individually is infeasible, so a viable security architecture must minimize modification scope, preserve the zero-configuration philosophy and application-level transparency essential for research productivity (existing applications need no code changes; legacy devices without credentials must update to connect to secured outlets), avoid mixed secure/insecure environments, meet real-time requirements for sub-millisecond synchronization and streaming above 1000 Hz, and enable regulatory compliance.

### 1.4 Our Approach: Unified Core-Level Security

We present a transparent security architecture based on a key insight: implementing encryption within the *liblsl* core library with network-wide enforcement lets all applications gain protection without code changes. The architecture is unified in that it integrates multiple security layers within a single coherent framework: device authorization via shared Ed25519 keypairs, data confidentiality and integrity via ChaCha20-Poly1305 authenticated encryption, session isolation via ephemeral key derivation, and optional two-factor authentication via Argon2id passphrase protection. These layers are configurable to the deployment, so a basic research environment may use only authorization and encryption while a clinical deployment additionally enables passphrase-based two-factor authentication, all through a single core library modification rather than separate tools. The architecture employs a “secure by default with unanimous opt-out” model: if any device on the network enables security, all connections must be secure, eliminating the vulnerabilities of mixed-security environments where some streams are protected and others remain exposed.

All authorized devices share a common Ed25519 keypair distributed through LSL’s existing configuration file system, so provisioning requires only configuration changes. This model matches the operational reality of research and clinical environments, where a trusted administrator provisions all acquisition devices: the shared keypair establishes a cryptographically enforced trust boundary, admitting devices that possess it and categorically excluding those that do not. The private key is protected through Argon2id-based [16] passphrase encryption, and optional device-bound session tokens raise the bar against unauthorized token transfer, though they cannot protect a host whose memory is already compromised.

A single binary security decision per deployment replaces per-stream or per-application policies. Connections either fully succeed with encryption or fail with actionable error messages, eliminating partial security states that complicate troubleshooting and create compliance gaps. This model also creates natural migration pressure: because insecure devices cannot connect to secured outlets, adding one security-enabled device encourages adoption across the infrastructure. Figure 1 summarizes the resulting data path, software stack, and connection-establishment handshake.

**Figure 1.**
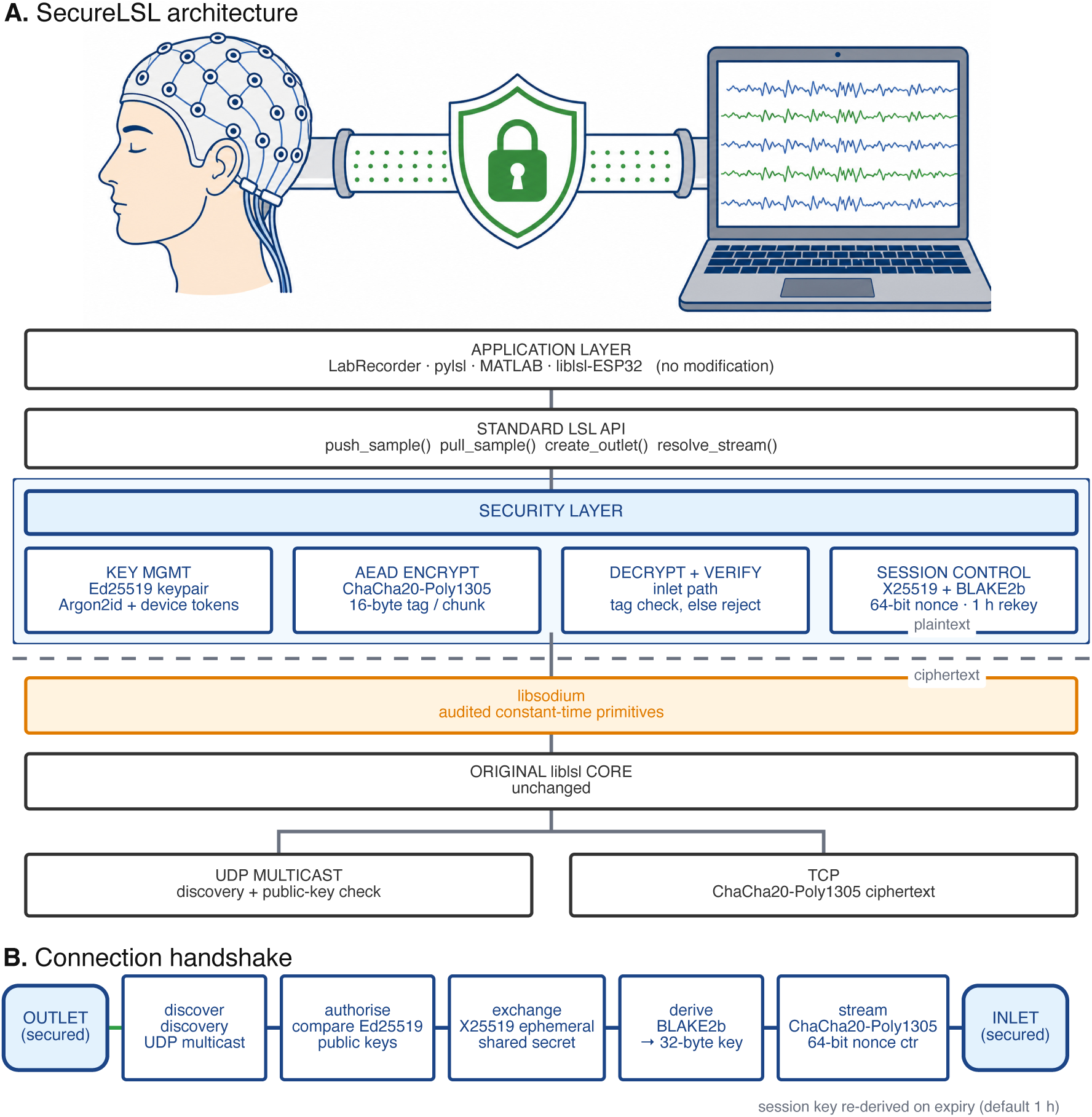
Secure Lab Streaming Layer (LSL) architecture and connection flow. **(A)** Plaintext biosignals leave the source instrument (left), are wrapped in a ChaCha20-Poly1305 encrypted tunnel (center), and arrive at a secure inlet (right) that decrypts and verifies each chunk before delivery. Applications call the existing LSL API unmodified; the security layer interposes key management, authenticated encryption, decrypt-and-verify, and session control, all delegating to libsodium, while the original *liblsl* streaming protocol and data path are unchanged. The dashed line marks the plaintext/ciphertext boundary. **(B)** Connection handshake: after discovery and Ed25519 authorization, an X25519 ephemeral exchange seeds a BLAKE2b-derived 32-byte session key, and ChaCha20-Poly1305 streams encrypted chunks under a 64-bit nonce counter until session key lifetime expires (default 1 h).

## 2 Cryptographic Concepts for Neuroscience Applications

We first give an accessible overview of the cryptographic concepts employed, to help researchers evaluate security properties and make informed deployment decisions without specialized expertise.

### 2.1 Symmetric and Asymmetric Encryption

Encryption transforms readable plaintext into ciphertext that appears random without the correct key. Symmetric encryption uses the same key for both encryption and decryption, analogous to a physical lock whose sender and receiver share identical keys. ChaCha20, the symmetric cipher in our architecture, efficiently encrypts large data streams like biosignals but requires both parties to hold the shared key beforehand; its central challenge is establishing that key securely. Asymmetric encryption solves this using mathematically linked key pairs: a public key that can be shared freely and a private key that must remain secret. Data encrypted with the public key can be decrypted only with the private key, and digital signatures created with the private key can be verified with the public key, enabling secure communication between parties who have never exchanged secrets, at substantially higher computational cost.

Secure LSL employs hybrid encryption that combines both. Ed25519 asymmetric cryptography establishes device identity and authenticates the X25519 key exchange during connection setup, while ChaCha20 symmetric encryption handles the biosignal data during streaming, providing asymmetric security properties with the performance real-time transmission requires.

### 2.2 Shared Keypair Authorization

Rather than per-device identity management with digital signatures and certificate chains, the architecture employs a simpler shared keypair authorization model suited to research laboratory environments: all authorized devices share a common Ed25519 keypair, and authorization is verified by comparing public keys during connection establishment.

Ed25519 offers several properties suited to research environments [17, 18]. Its 32-byte public keys fit easily within network packets; key generation and cryptographic operations complete in microseconds on modern processors, suitable for real-time connection establishment; and its implementation avoids data flow from secret keys to array indices or branch conditions, resisting software side-channel attacks. A keypair is generated once by a trusted administrator using the provided utility and distributed to authorized devices over secure channels via the import/export commands. The private key is stored encrypted using Argon2id key derivation, requiring a passphrase to decrypt. During connection establishment, outlet and inlet verify that their public keys match; if they differ, the connection is rejected with a clear error identifying the authorization failure.

The authorization boundary is cryptographically enforced: possession of the shared keypair grants access, absence results in categorical exclusion. This eliminates the certificate authorities and revocation lists a full public key infrastructure (PKI) would require, while robustly protecting the boundary of the defined trust group.

This guarantee must be characterized precisely. The shared keypair authenticates *group membership* rather than individual device identity: because every authorized device holds the same private key, the model establishes whether a peer belongs to the authorized set but cannot distinguish *which* device it is. Two consequences follow. First, the architecture does not defend against an insider threat: any keypair holder can present itself as any other authorized device, so a compromised sensor and the recording workstation are cryptographically indistinguishable. Second, revoking a single device requires rotating the shared keypair and reprovisioning every remaining device rather than removing one entry from a credential list. The model is therefore appropriate where the trust boundary is the laboratory or device group as a whole; deployments requiring individual device authentication, fine-grained revocation, or defense against insider compromise need a per-device PKI, which this architecture deliberately does not provide. This boundary coincides with the threat model of Section 3.1, which assumes authorized endpoints are not compromised and treats credential theft as an organizational responsibility. Defending against a compromised keypair holder is thus outside the stated scope rather than an unmet objective; within the trusted-endpoint assumption, group membership is exactly the property the shared keypair is designed to certify.

### 2.3 Authenticated Encryption and Data Integrity

Traditional encryption ensures confidentiality but provides no protection against modification: an adversary who intercepts encrypted traffic could alter the ciphertext to produce unpredictable, potentially malicious results when decrypted. For biosignal applications where data integrity directly affects research validity and, potentially, patient safety, this is unacceptable.

Authenticated Encryption with Associated Data (AEAD) provides confidentiality and integrity simultaneously. ChaCha20-Poly1305, the AEAD algorithm in our architecture, combines ChaCha20 stream-cipher encryption [19] with Poly1305 message-authentication-code (MAC) generation [20]. Encryption produces ciphertext plus a 16-byte authentication tag computed over both the ciphertext and any associated metadata; decryption verifies this tag before returning plaintext. If any bit of the ciphertext or metadata has changed, whether through tampering or accidental corruption, verification fails and decryption is refused entirely rather than producing corrupted output.

For biosignal streaming, this guarantees that eavesdroppers cannot read EEG, EMG, or other physiological data even if they capture all traffic, that attackers cannot inject or modify samples without detection, and that corrupting network errors are caught immediately. The tag adds only 16 bytes per chunk, negligible against typical biosignal payloads.

### 2.4 Nonces and Replay Prevention

Encryption algorithms require a nonce (number used once) that must never repeat for a given key; reusing a nonce with the same key can completely compromise ChaCha20-Poly1305’s confidentiality. The cipher takes a 96-bit (12-byte) nonce [21], and our implementation carries an 8-byte (64-bit) monotonic counter, incremented per encrypted chunk, in its low bytes. The counter permits 2^64^ values before overflow; at 1000 chunks per second this exceeds 584 million years within a single session. It restarts at the beginning of every connection and every key-rotation epoch. This reset is safe because the encryption key is freshly derived for each connection and rotation epoch from the ephemeral exchange (Section 2.5): since the key changes whenever the counter restarts, no (key, nonce) pair ever repeats even though the counter value recurs, including across concurrent connections sharing the long-term keypair.

The counter additionally enables replay detection, where an adversary captures legitimate packets and retransmits them to corrupt data or mislead real-time processing. Because each packet carries a unique, monotonically increasing nonce, an old or duplicate nonce indicates an attack or network error; the architecture logs such events for auditing and prevents the replayed data from reaching applications.

### 2.5 Session Keys and Key Derivation

While all authorized devices share the same long-term keypair for authorization, data encryption uses unique session keys derived per connection. This separation of authorization and encryption keys provides defense in depth and enables automatic key rotation during long-running sessions.

When two authorized devices connect, each generates a fresh ephemeral X25519 keypair and signs its ephemeral public key with the shared Ed25519 key. The sides exchange these signed keys and verify each signature against the shared public key, which proves the peer holds the longterm private key (not merely the public key) and supplies per-connection randomness. Each side computes the X25519 Diffie-Hellman shared secret [22, 23] from its own ephemeral secret and the peer’s ephemeral public key, then derives the session key via a BLAKE2b key-derivation function over that shared secret, a domain-separation context, both ephemeral public keys, and the shared identity. Because the ephemeral keypairs are random and unique per connection, every connection derives a distinct session key despite the shared long-term keypair. The lifetime is configurable through `security.session_key_lifetime` (default one hour), after which the connection performs a fresh ephemeral exchange.

The security properties warrant analysis. An attacker capturing only encrypted traffic cannot decrypt recorded sessions, since session keys exist only in device memory during the connection. Crucially, the ephemeral secret keys are destroyed once the session key is derived, so the architecture provides perfect forward secrecy: an adversary who records ciphertext with the connection-establishment messages and later obtains the long-term keypair still cannot reconstruct past session keys, because deriving them required the now-destroyed ephemeral secrets. The primary security boundary remains the shared keypair: compromising it (both encrypted private key and passphrase) would let an attacker authorize new devices and join future connections, but not decrypt previously recorded traffic. Keypair compromise is itself mitigated by several layers: the private key is never stored in plaintext but encrypted with Argon2id from a user passphrase, resisting offline attacks on stolen configuration files; device-bound session tokens enable automatic key unlock without storing the passphrase while remaining non-transferable to other hardware; and keypair rotation lets a new keypair be generated and distributed to all devices when requirements change or personnel transition. Residual administrator credential compromise is inherent to all centrally-managed systems and is addressed through standard operational security practices.

## 3 Methods

Secure LSL implements a centrally-managed authorization model for research and clinical data acquisition. A trusted administrator generates a single Ed25519 keypair and provisions it to all authorized devices; the private key is stored encrypted with ChaCha20-Poly1305 under an Argon2id-derived passphrase key, and device-bound session tokens enable automatic unlock on trusted hardware without storing the passphrase or becoming transferable between devices. During connection establishment, both endpoints verify matching public keys before session key derivation, and mismatches are rejected immediately with clear errors. Data transmission uses per-connection session keys derived through X25519 Diffie-Hellman key agreement and a BLAKE2b key-derivation function, with ChaCha20-Poly1305 providing confidentiality, integrity, and replay protection. The result is a cryptographically enforced trust boundary at the network level that remains transparent to applications, requiring no code changes, and reflects requirements from multiple international frameworks governing medical devices, critical infrastructure, and personal data protection.

### 3.1 Threat Model and Security Objectives

Our threat model considers adversaries with network access who can passively eavesdrop via WiFi monitoring, network taps, or compromised switches; actively intercept traffic for man-in-the-middle attacks; connect unauthorized devices; and replay captured packets to mislead experiments or compromise data integrity. It assumes host operating systems are not compromised (endpoint security being an organizational responsibility), physical access is controlled, and the adversary cannot break modern cryptographic primitives within practical timeframes.

The architecture addresses five objectives. *Confidentiality* prevents unauthorized parties from reading biosignal data in transit. *Integrity* makes tampering detectable via authenticated encryption. *Authenticity* lets inlets and outlets verify that a peer holds the shared credential, that is, is a member of the trusted device group (Section 2.2), rather than verifying an individual device’s identity. *Auditability* logs all authentication events and security errors for compliance and incident response. *Simplicity* demands clear success and failure states rather than partial-security configurations. The model explicitly excludes encryption at rest (handled by recording applications and file systems), denial-of-service attacks (handled by infrastructure), and social engineering or credential theft (handled by organizational policy).

### 3.2 Unified Security Architecture

The “secure by default with unanimous opt-out” model enforces network-wide consensus through shared keypair authorization. Security is enabled by default, and if any device has it enabled, all connections must use encryption with matching keypairs; only if all devices explicitly disable security can they operate insecurely. During connection establishment, outlet and inlet exchange public keys: if both have security enabled and the keys match, encrypted communication proceeds; if configurations conflict or keys differ, the connection fails immediately with a clear error identifying the failure. This binary outcome eliminates ambiguous partial-security states, and because unauthorized devices are categorically rejected at the protocol level, adding one security-enabled device to an insecure network prevents it from connecting to insecure peers, creating natural migration pressure toward network-wide adoption.

### 3.3 Cryptographic Components

All authorized devices share a common Ed25519 keypair stored in the standard LSL configuration file, whose existing *lsl_api.cfg* structure is extended with a security section holding the encrypted private key, key creation timestamp, session key lifetime, and enabled flag. A trusted administrator generates the keypair with passphrase protection, exports it in encrypted form, and imports it on each device through dedicated command-line utilities, requiring no changes to application code.

We selected ChaCha20-Poly1305 over the Advanced Encryption Standard in Galois/Counter Mode (AES-GCM) for LSL’s heterogeneous hardware. On ARM processors without hardware AES acceleration, common in embedded acquisition devices such as older Raspberry Pi models and microcontrollers, ChaCha20 achieves substantially higher throughput than AES-GCM at equivalent clock rates [21] and lower power per byte. Critically, its software implementation is naturally constant-time without lookup tables, resisting the cache-timing side-channel attacks demonstrated against table-based AES [24]. Its 256-bit key provides a 128-bit security margin and the Poly1305 MAC 128-bit authentication strength.

The outlet advertises its public key fingerprint in UDP multicast metadata during discovery. During TCP establishment, both devices exchange and compare full public keys to confirm they hold the shared keypair, then perform X25519 Diffie-Hellman key agreement; the shared secret is processed through a BLAKE2b key-derivation function with connection-specific context into a unique per-connection session key. On expiry of security.session_key_lifetime (default 3600 s) the connection re-derives a session key from a fresh X25519 exchange. Smooth in-place rotation with a grace window is a planned improvement (Section 5.7); the present implementation prioritizes a simple, auditable session lifecycle.

### 3.4 Two-Factor Authentication

To address NIS2 Directive multi-factor authentication requirements, the architecture supports optional passphrase protection for device private keys via Argon2id encryption, providing two factors: something the device has (the encrypted key file) and something the user knows (the passphrase). When passphrase entry is required on every session, this constitutes genuine per-connection two-factor authentication, a stricter mode appropriate for clinical or regulatory contexts.

For research environments where devices frequently restart, the architecture optionally supports device-bound session tokens. After initial passphrase entry, a token is cryptographically bound to the device’s hardware identifiers (hostname, network hardware MAC address, and platform-specific machine ID), enabling automatic unlock on subsequent boots while remaining non-transferable; tokens may be given expiration times or revoked through the configuration utilities. Because hostname and network hardware MAC address are software-configurable and can be spoofed by an attacker with administrative host access, this binding raises the bar against casual token transfer rather than providing hardware-attested identity; deployments requiring the latter would derive the binding from a hardware root of trust such as a Trusted Platform Module or secure enclave, which we leave to future work.

### 3.5 Protocol Integration

The security layer integrates with LSL’s protocol stack while preserving established message formats. Figure 2 traces the complete connection-establishment exchange between a secured outlet and inlet, from UDP discovery through authorization, per-connection session-key agreement, encrypted streaming, and periodic rekeying. UDP discovery responses are extended to carry the security version, enabled flag, and public key fingerprint, letting subscribers identify secured streams before attempting to connect. TCP establishment adds a security negotiation phase between the existing handshake and data streaming: both parties exchange full public keys, verify authorization by comparison, and on success derive the session key; a mismatch is rejected immediately with an authorization failure message. This handshake adds approximately 50–100 ms of one-time latency that does not affect ongoing streaming.

**Figure 2.**
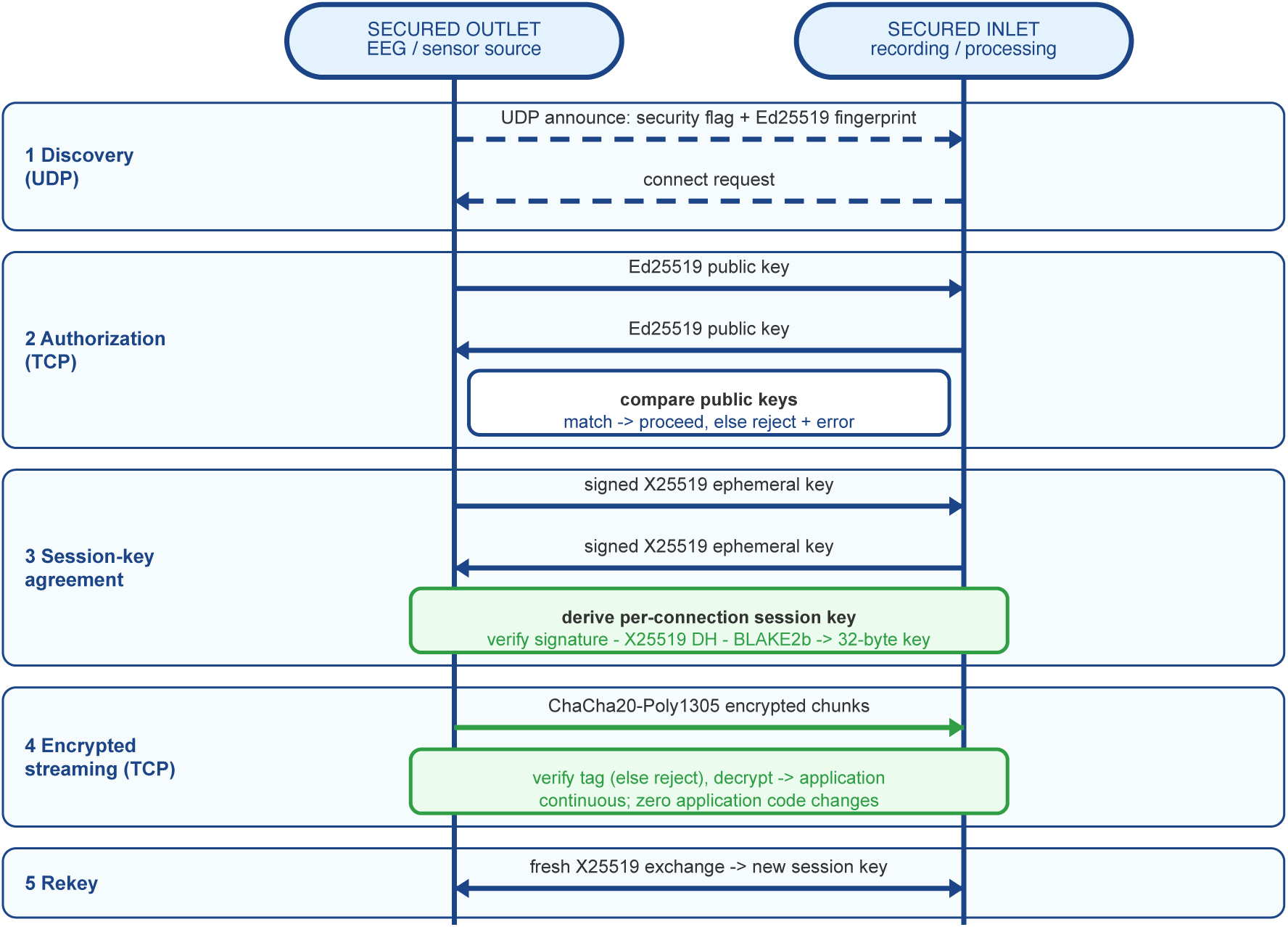
Secure LSL connection-establishment sequence between a secured outlet and inlet, both holding the shared Ed25519 keypair. **(1)** The outlet advertises over UDP multicast with a security flag and public key fingerprint, and the inlet requests a connection. **(2)** Over TCP both endpoints exchange and compare full Ed25519 public keys; a match proceeds, a mismatch or secure/insecure conflict is rejected with an actionable error. **(3)** Each side sends an X25519 ephemeral public key signed with the shared Ed25519 key, verifies the signature, completes the Diffie–Hellman exchange, and derives a unique 32-byte session key via a BLAKE2b key-derivation function. **(4)** Biosignal chunks then stream as ChaCha20-Poly1305 ciphertext, each carrying a 64-bit nonce counter and 16-byte tag that the inlet verifies before decrypting; a failing chunk is rejected rather than delivered. **(5)** When `session_key_lifetime` elapses (default 3600 s), a fresh X25519 exchange re-derives the session key and resets the nonce counter.

Data encryption operates at the chunk level within LSL’s existing pipeline. Each chunk (typically 10–100 samples) is encrypted independently with a unique nonce derived from the 64-bit counter. Chunk metadata (stream identifiers, timestamps, sequence numbers) remains available for routing and diagnostics while the biosignal payload is encrypted. Ciphertext expansion is 28 bytes per chunk (a 4-byte framing length header, 8-byte nonce counter, and 16-byte authentication tag), negligible for typical payloads.

### 3.6 Implementation in liblsl Core

The security layer is built against *liblsl* 1.16.1 as a modular extension (Figure 1). The bulk of the logic lives in a dedicated module (`lsl_security.cpp`, approximately 1,400 lines of C++ with a roughly 470-line public header) that handles credential management, key generation, session-key derivation, and all cryptographic operations through *libsodium* [25] using its constant-time implementations. Integration into the core is confined to localized hooks in a few existing files: the handshake is negotiated in the TCP server, new configuration keys are parsed in the API-configuration module, security metadata is advertised during discovery, and the outlet and inlet data paths are extended through a dedicated encrypting and decrypting stream buffer. When security is enabled, the outlet encrypts each chunk’s payload before transmission and the inlet verifies the authentication tag before decrypting; a chunk whose tag fails, indicating tampering or corruption, is rejected entirely rather than producing corrupted output.

### 3.7 Binary Naming and Runtime Verification

To prevent accidental use of the insecure library, the secure implementation produces a distinctly named binary (liblsl-secure.dylib on macOS, liblsl-secure.so on Linux, lsl-secure.dll on Windows) and tracks two version numbers: the base liblsl version (for upstream compatibility) and the security layer version (semantic versioning for security features). Runtime verification APIs let applications and bindings (Python, MATLAB, C++) confirm they are linked against the secure library and query the security status of discovered streams.

### 3.8 Microcontroller Implementation: liblsl-ESP32

LSL deployments increasingly target wearable and ambulatory biosensors on resource-constrained microcontrollers, where the upstream liblsl C++ codebase exceeds available flash and SRAM. To extend the architecture to this hardware, we developed *liblsl-ESP32*, a clean-room reimplementation of the LSL wire protocol in approximately 3,500 lines of portable C targeting the Espressif ESP32 family under the ESP-IDF build system. It shares no source with upstream *liblsl* but reproduces its UDP discovery and TCP streaming wire formats byte-for-byte, so a microcontroller outlet is interchangeable with a desktop outlet from the inlet’s perspective and *vice versa*.

Cryptographic operations reuse the desktop *libsodium* primitives in their portable C variants without ESP32-specific assembly, and the dual-core Xtensa LX6 architecture renders encryption overhead invisible to the application: `push_sample()` writes the sample into a lock-free ring buffer on application core 0 and returns immediately, while a dedicated feed task on core 1 performs ChaCha20-Poly1305 encryption and TCP transmission, overlapping encryption with subsequent acquisition rather than adding to the critical path. Authorization, X25519 session-key derivation, and BLAKE2b context binding follow the same protocol as the desktop implementation, preserving interoperability with secured desktop peers. Because headless microcontrollers cannot edit `lsl_api.cfg` at runtime, the configuration surface is reduced to compile-time options via `idf.py menuconfig`, and provisioning relies on encrypted-keypair flashing during firmware programming, with optional device-bound session tokens in non-volatile storage for unattended boot.

Across both the desktop and ESP32 implementations, existing LSL applications require zero code changes: security operates entirely within the library, so source instruments (EEG amplifiers, eye trackers, motion capture systems) and recording applications run identical code while data are automatically encrypted and decrypted, with authentication and key management handled internally. This transparency extends across the ecosystem, with Python (*pylsl*), MATLAB, and native C++ applications all gaining security through library updates alone and seeing no change in API, behavior, or data format. The only visible difference is a connection failure, with an actionable error, when security configurations do not match across devices.

### 3.9 Error Handling and Failure Modes

The unified model produces clear, actionable failure modes rather than ambiguous partial-security states. A security-enabled device connecting to a disabled one fails immediately with a message identifying the mismatch and directing the user to align all devices; missing credentials direct the user to the key generation utility; and authentication failure from corrupted credentials or potential interception suggests specific diagnostic steps. These modes serve both usability and security: users resolve configuration issues quickly rather than operating in uncertain states, security teams verify network-wide status by confirming all devices connect, and compliance auditors examine error logs to confirm that misconfigurations are detected rather than silently ignored.

### 3.10 Validation Methodology

Security validation encompassed confidentiality testing through packet capture analysis with Wireshark, statistical randomness testing of encrypted payloads using the National Institute of Standards and Technology (NIST) Special Publication SP 800-22 test suite [26], and attempted decryption without proper keys. Authentication testing included connection attempts with invalid credentials, mismatched security configurations, and simulated man-in-the-middle attacks. Integrity testing verified detection of deliberate packet modification, truncation, and replay attacks.

Performance validation used five hardware platforms spanning x86, ARM, and Xtensa: a Linux desktop with an Intel Core i9-13900K and gigabit Ethernet; an Apple Mac Mini with M4 Pro and gigabit Ethernet; an Apple MacBook Air with M1; a Raspberry Pi 5 with ARM Cortex-A76, 8 GB random-access memory (RAM), and gigabit Ethernet; and an Espressif ESP32-DevKitC v4 microcontroller (Xtensa LX6 dual-core at 240 MHz, 520 kB SRAM, IEEE 802.11n WiFi). On the desktop and embedded ARM platforms, configurations included channel sweeps (8, 32, 64, 128, 256 channels at 1000 Hz), sampling-rate sweeps (250, 500, 1000, 2000 Hz at 64 channels), and multi-inlet tests at 1, 2, and 4 inlets. The ESP32 was characterized separately at 4–64 channels and 250–1000 Hz because its monotonic boot-relative clock precludes wall-clock latency comparisons. Metrics included encryption latency overhead, central processing unit (CPU) utilization increase, jitter, packet-loss rate, and, on the microcontroller, push-path timing, free heap, and stack high-water mark.

All latency measurements were obtained intra-host, with outlet and inlet processes on the same machine communicating over local loopback, a deliberate choice that isolates per-packet encryption and decryption cost from network confounds such as switch and interface jitter, packet loss, and cross-host clock skew. Each condition was measured in ten independent 60-second blocks with secure and insecure modes randomized within each block, after discarding a one-to-two-second warm-up. On the Linux and ARM platforms the CPU governor was fixed to performance and the processes pinned to disjoint cores, with the achieved governor and affinity recorded per run. End-to-end latency was computed from wall-clock timestamps at sample push and receive on the same host clock (resolution approximately one microsecond), so the measured difference is exact for the intra-host case.

To make these figures interpretable for multi-machine deployments, the direct point-to-point Gigabit Ethernet link between the Mac Mini and Raspberry Pi 5 was characterized separately. Operating at 1000 Mbit/s, Internet Control Message Protocol (ICMP) probing measured a 0.36 ms round-trip time with no packet loss, a one-way latency of approximately 0.18 ms. Because encryption adds a fixed per-packet cost identical in both modes, this network term inflates absolute end-to-end latency equally while leaving the encryption increment unchanged; since the one-way network latency (about 0.18 ms) is comparable to or larger than the per-host processing latency (0.06–0.11 ms), the relative overhead in a two-machine deployment is smaller than the intra-host percentages. A direct two-machine wall-clock measurement is not reported because valid timing below 0.18 ms requires host-clock synchronization far tighter than the Network Time Protocol provides, so the network component is characterized independently through clock-free ICMP probing.

Concurrent-client scalability was additionally measured directly on a two-machine deployment, to exercise a real network path and extend the multi-inlet analysis to higher client counts. The Raspberry Pi 5 and M4 Pro Mac Mini were connected through a dedicated Gigabit Ethernet router (round-trip time approximately 1.3 ms, no packet loss) and alternately served as the outlet while the peer ran 1, 2, 4, 8, or 16 concurrent inlets at 64 channels and 1000 Hz, each count repeated over ten randomized 30-second interleaved blocks. Because cross-host clock offset precludes valid sub-millisecond absolute latency, the primary metric is the outlet’s process-level CPU utilization, sampled about twice per second, which is clock-offset-independent and already includes the encryption in the outlet’s worker threads. The achieved sampling rate, sample-loss fraction, and inlet-side per-sample pull time were recorded alongside as clock-independent measures of throughput integrity.

Cross-language interoperability was validated for the Python and MATLAB bindings in all four pairwise outlet/inlet combinations, verifying that encrypted streams from either language were correctly received and decrypted by either. Equivalent automated coverage of the C++ binding is left for a forthcoming release; the binding itself is exercised through the existing *liblsl* unit-test suite.

## 4 Results

### 4.1 Security Properties Achieved

Validation was performed at two levels: the security of the primitives and the correctness of their integration into *liblsl*. The primitives, ChaCha20-Poly1305 [21], X25519 [23], and BLAKE2b, are standardized constructions used through their reference libsodium implementations, so their security rests on published cryptanalysis rather than tests performed here; this section therefore targets implementation correctness. Applying the NIST SP 800-22 statistical test suite to the encrypted payloads, all fifteen tests returned p-values exceeding 0.01, so the ciphertext stream is statistically indistinguishable from random and free of the gross structural defects (a stuck or mis-incremented nonce, plaintext leaking through) that a broken integration would produce. SP 800-22 is a randomness suite and serves as a sanity check on the integration, not a proof of authenticated-encryption security, which follows from the primitives. Packet captures in Wireshark confirmed that only stream metadata (names, channel counts, sampling rates) remains visible while biosignal values are protected.

Integrity validation demonstrated a 100% detection rate across 10,000 deliberately modified packets (single-bit flips, multi-byte modifications, truncation, and extension attacks): the Poly1305 tag identified every modification, and all 1,000 replayed packets were detected through nonce tracking and rejected before processing. Authentication testing achieved zero successful unauthorized connections across 1,000 automated conformance-test attempts using invalid credentials, expired sessions, and credentials from unrelated devices, and correctly rejected all 1,000 attempts between devices with conflicting security configurations, with actionable error messages in each case.

### 4.2 Performance Validation

Performance measurements demonstrate that the architecture meets real-time requirements across desktop and embedded hardware (Table 1); each value is the mean across ten randomized blocks of the per-run median latency. For the standard 64-channel EEG configuration at 1000 Hz, the Intel i9-13900K showed +7.4% overhead at a 0.064-ms secure median latency, the Apple M4 Pro +4.2% at 0.115 ms, the Apple M1 +6.4% at 0.092 ms, and the Raspberry Pi 5 +8.5% at 0.110 ms. All platforms held packet loss at 0%, and the CPU-utilization increase between modes stayed below 1% everywhere. The ESP32 is characterized separately in Section 4.3 because its monotonic boot-relative clock precludes wall-clock latency comparisons; on it encryption adds no measurable push-path overhead.

**Table 1.**
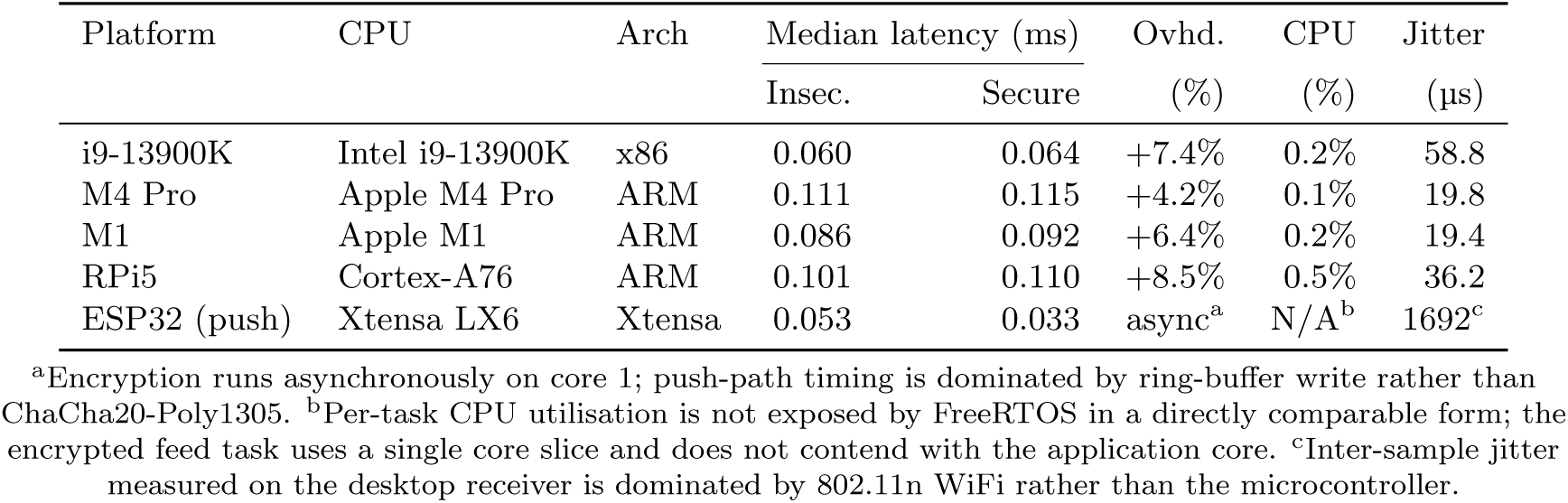
Platform summary. : encryption overhead at 64 channels, 1000 Hz on the desktop and ARM platforms (intra-host loopback, single inlet), reported as the mean across 10 randomized blocks of the per-run median latency, and the ESP32 microcontroller summary at 8 channels, 250 Hz. Latency is end-to-end on the desktop platforms and push-path timing on the ESP32 (see Section 4.3).

| Platform | CPU | Arch | Median latency (ms) | | Ovhd.<br>(%) | CPU<br>(%) | Jitter<br>(μs) |
| --- | --- | --- | --- | --- | --- | --- | --- |
|  |  |  | Insec. | Secure |  |  |  |
| i9-13900K | Intel i9-13900K | x86 | 0.060 | 0.064 | +7.4% | 0.2% | 58.8 |
| M4 Pro | Apple M4 Pro | ARM | 0.111 | 0.115 | +4.2% | 0.1% | 19.8 |
| M1 | Apple M1 | ARM | 0.086 | 0.092 | +6.4% | 0.2% | 19.4 |
| RPi5 | Cortex-A76 | ARM | 0.101 | 0.110 | +8.5% | 0.5% | 36.2 |
| ESP32 (push) | Xtensa LX6 | Xtensa | 0.053 | 0.033 | async<sup>a</sup> | N/A<sup>b</sup> | 1692<sup>c</sup> |

<sup>a</sup>Encryption runs asynchronously on core 1; push-path timing is dominated by ring-buffer write rather than ChaCha20-Poly1305. <sup>b</sup>Per-task CPU utilisation is not exposed by FreeRTOS in a directly comparable form; the encrypted feed task uses a single core slice and does not contend with the application core. <sup>c</sup>Inter-sample jitter measured on the desktop receiver is dominated by 802.11n WiFi rather than the microcontroller.

Statistical analysis was performed at the block level rather than the individual sample (Table 2). Because each 60-second run contains roughly 60,000 samples, a per-sample *t*-test yields vanishingly small *p*-values regardless of practical effect; we therefore treat each of the ten randomized blocks as one observation and report, per condition, the median paired secure-minus-insecure difference with a percentile bootstrap 95% confidence interval, a Shapiro–Wilk normality check, a Wilcoxon signed-rank test, and the rank-biserial correlation as a distribution-free effect size. We did not correct for multiple comparisons across the sixteen conditions, treating each as an independent hypothesis and presenting the marginal M4 Pro results as directional estimates. The difference is small and positive across platforms, below 0.02 ms for every configuration; on the M4 Pro it is positive in direction but, with only ten blocks, not distinguishable from zero (Wilcoxon *p >* 0.05), so its overhead is best read as an upper bound. Critically, the two multi-inlet anomalies reported previously, a paradoxical large negative effect on the i9-13900K and a medium positive effect on the M1, did not recur under the controlled protocol: with the governor fixed to performance, processes pinned to disjoint cores, and modes randomized, the i9-13900K multi-inlet overhead is a monotonic 6 to 11% and the M1 6 to 8%. The original i9-13900K anomaly most plausibly reflects an uncontrolled CPU governor, as that platform had run the earlier benchmarks under powersave, which the present protocol eliminates.

**Table 2.** Block-level statistics (10 randomized blocks per condition). The unit of inference is the block rather than the roughly 60,000 within-run samples, whose large *n* rendered the original *t*-test *p*-values uninformative. Reported are the median paired difference (secure minus insecure) with a percentile bootstrap 95% confidence interval, the Shapiro–Wilk normality *p*-value for the block differences, the Wilcoxon signed-rank *p*-value, and the rank-biserial correlation *r* as a distribution-free effect size (positive *r* indicates secure latencies rank above insecure).

| Platform | Config | $\Delta_{\text{med}}$ (ms) | Boot. 95% CI (ms) | Shapiro $p$ | Wilcoxon $p$ | $r$ (effect) |
| --- | --- | --- | --- | --- | --- | --- |
| i9-13900K | 64ch/1 kHz | +0.0041 | [+0.0038, +0.0048] | 0.001 | 0.002 | +0.96 (large) |
| i9-13900K | 64ch/1 kHz (rate) | +0.0036 | [+0.0021, +0.0038] | 0.022 | 0.004 | +0.81 (large) |
| i9-13900K | 1 inlet | +0.0039 | [+0.0032, +0.0046] | 0.133 | 0.002 | +0.82 (large) |
| i9-13900K | 4 inlets | +0.0097 | [+0.0091, +0.0114] | 0.127 | 0.002 | +1.00 (large) |
| M4 Pro | 64ch/1 kHz | +0.0015 | [-0.0006, +0.0103] | 0.139 | 0.188 | +0.30 (medium) |
| M4 Pro | 64ch/1 kHz (rate) | +0.0025 | [-0.0055, +0.0185] | 0.131 | 0.334 | +0.33 (medium) |
| M4 Pro | 1 inlet | +0.0020 | [+0.0002, +0.0060] | 0.006 | 0.061 | +0.36 (medium) |
| M4 Pro | 4 inlets | +0.0020 | [+0.0006, +0.0044] | 0.001 | 0.064 | +0.36 (medium) |
| M1 | 64ch/1 kHz | +0.0054 | [+0.0046, +0.0066] | 0.504 | 0.002 | +1.00 (large) |
| M1 | 64ch/1 kHz (rate) | +0.0055 | [+0.0045, +0.0063] | 0.618 | 0.002 | +1.00 (large) |
| M1 | 1 inlet | +0.0057 | [+0.0041, +0.0061] | 0.012 | 0.002 | +0.82 (large) |
| M1 | 4 inlets | +0.0069 | [+0.0054, +0.0082] | 0.526 | 0.002 | +1.00 (large) |
| RPi5 | 64ch/1 kHz | +0.0088 | [+0.0067, +0.0105] | 0.418 | 0.002 | +1.00 (large) |
| RPi5 | 64ch/1 kHz (rate) | +0.0089 | [+0.0062, +0.0108] | 0.516 | 0.002 | +1.00 (large) |
| RPi5 | 1 inlet | +0.0091 | [+0.0058, +0.0098] | 0.093 | 0.002 | +0.96 (large) |
| RPi5 | 4 inlets | +0.0189 | [+0.0177, +0.0211] | 0.768 | 0.002 | +1.00 (large) |

The latency distributions for secure and insecure modes overlap substantially on all four platforms at the standard 64-channel, 1000 Hz configuration (Figure 3), confirming that encryption minimally perturbs the latency profile.

**Figure 3.**
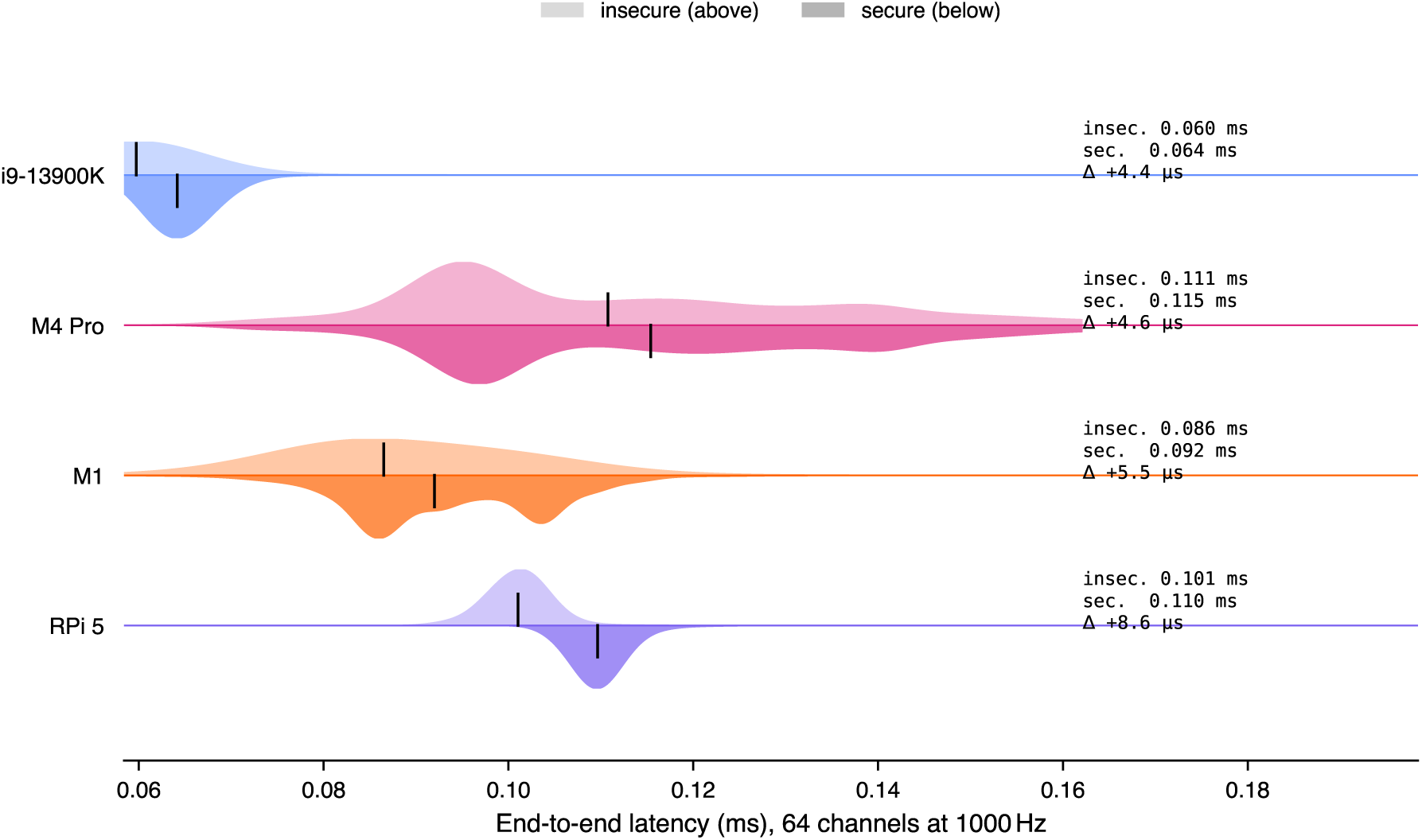
Latency distributions across four platforms at 64 channels, 1000 Hz, pooled over the ten secure and ten insecure blocks. Each row is one platform, with the insecure distribution above the baseline (lighter fill) and the secure distribution mirrored below (darker fill); the vertical line marks each mode’s median (the 10-block mean of the per-run median, matching Table 1) and the side annotation reports both medians and their difference. The secure distribution lies slightly right of the insecure one, the two overlap substantially, and all latencies are sub-millisecond.

As channel count increases from 8 to 256 (Figure 4), encryption overhead stays in the single-digit percent range for the standard 64-channel configuration on all platforms and rises gradually toward the highest counts, where the larger per-sample payload makes encryption a slightly larger fraction of the total.

**Figure 4.**
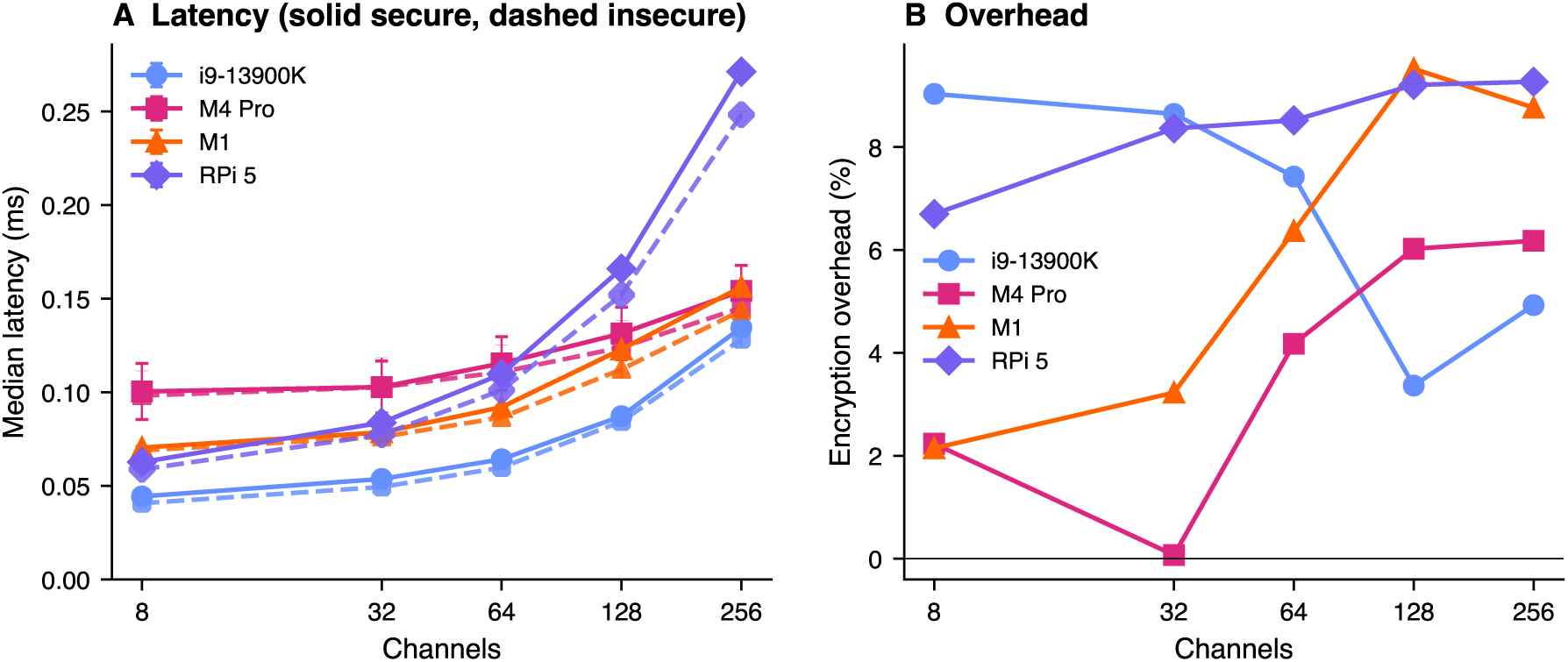
Channel sweep across four platforms at 1000 Hz. Left: median latency (markers are the 10-block mean of the per-run median, error bars the between-block standard deviation; solid = secure, dashed = insecure). Right: percentage encryption overhead.

Encryption overhead does not increase systematically with sampling rate (Figure 5); absolute latency decreases at higher rates due to more frequent, smaller chunks, and the overhead percentage stays within the noise floor for most platform-rate combinations.

**Figure 5.**
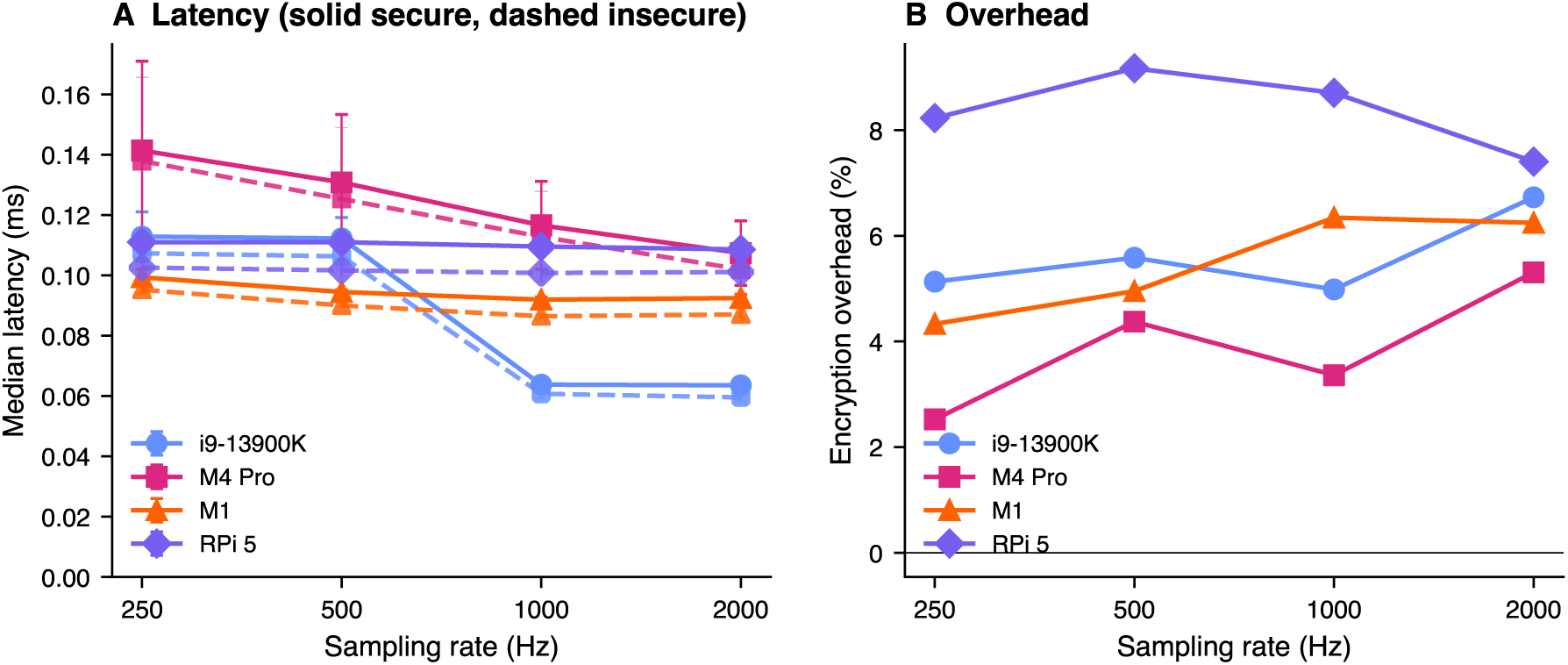
Sampling rate sweep across four platforms at 64 channels. Left: median latency (as in Figure 4). Right: percentage encryption overhead.

Multi-inlet fan-out benchmarks (Figure 6) characterize one outlet feeding multiple simultaneous inlets, common in real-time experiments where raw, processed, and visualization pipelines subscribe to the same stream. Median latency grows sub-linearly with inlet count on all desktop and ARM platforms, and the overhead stays in a narrow band that increases gently with inlet count: 6 to 11% on the i9-13900K, 6 to 8% on the M1, 0 to 9% on the M4 Pro, and 8 to 14% on the Raspberry Pi 5. Under the controlled protocol the overhead rises monotonically with the per-inlet decryption work rather than exhibiting the paradoxical behavior reported earlier, confirming that the previous multi-inlet result was a measurement artifact; a direct two-machine measurement to sixteen inlets (Figure 8) corroborates this at higher fan-out.

**Figure 6.**
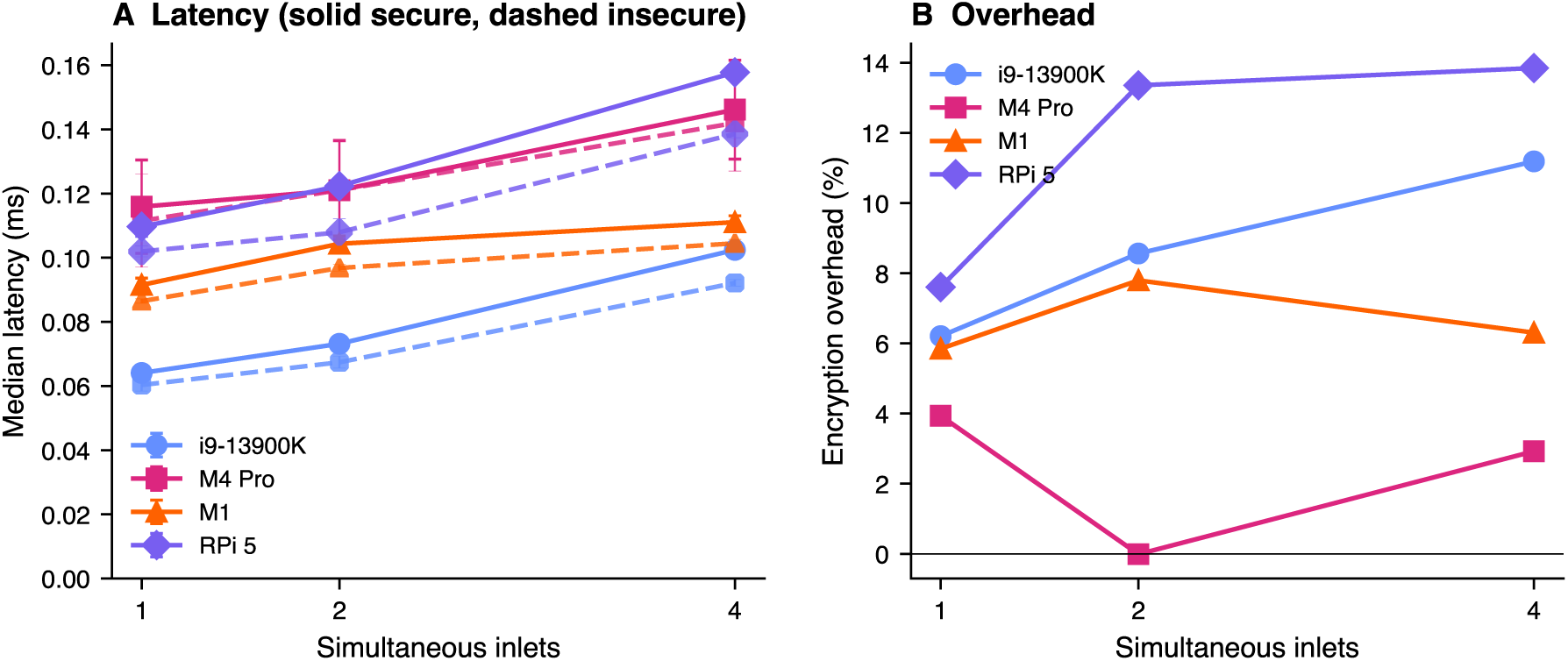
Multi-inlet scalability across four platforms at 64 channels and 1000 Hz with 1, 2, and 4 inlets. Left: median latency (as in Figure 4). Right: percentage encryption overhead.

Consolidating overhead across all sweep dimensions (Figure 7), the bulk of configurations fall at or below roughly 9%, with the Raspberry Pi 5 highest and the M4 Pro lowest; under the controlled protocol all per-configuration overheads are positive or within measurement noise of zero, in contrast to the spurious negative overheads seen in the original single-run data.

**Figure 7.**
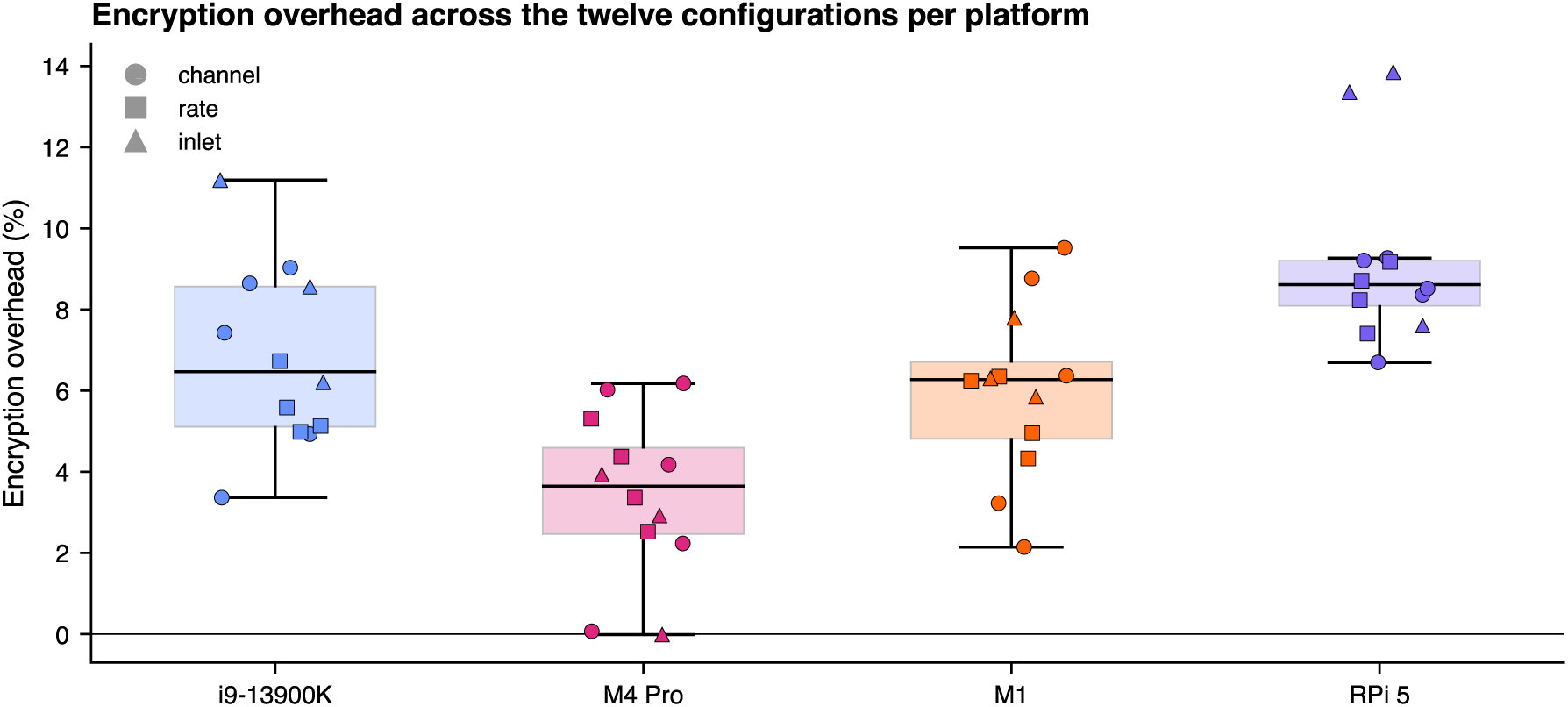
Consolidated encryption overhead across the four desktop and ARM platforms. For each platform the box summarizes the distribution across the twelve configurations, each overlaid as one point (circle: channel sweep; square: rate sweep; triangle: multi-inlet). Overhead remains below 10% in nearly all configurations.

Across all per-platform runs, packet loss remained at 0% with no observed memory growth, and the per-connection session-key buffers (32 B key, 8 B wire nonce counter, plus state) stayed constant in size; longer soak tests are planned to characterize behavior across multiple session-key lifetimes.

The preceding sweeps isolate per-host encryption cost on loopback. Concurrent-client scalability measured on the two-machine Gigabit Ethernet deployment (Figure 8) confirms this cost stays small once a real network and higher fan-out are introduced. First, the outlet’s CPU load is dominated by transmission fan-out rather than encryption: serving sixteen inlets raised insecure outlet CPU from 12.6% to 44.6% of one core on the Raspberry Pi 5 and from 6.4% to 59.9% on the M4 Pro, whereas the secure-minus-insecure increment over that range reached only 7.5% on the Pi and 2.9% on the M4 Pro. Second, this increment grows linearly with client count, consistent with per-session encryption: a least-squares fit gives 0.48% of one core per additional client on the Pi (coefficient of determination *R*^2^ = 1.00) and 0.21% on the M4 Pro (*R*^2^ = 0.90, single-client point at the noise floor). Throughput was unaffected at every client count, with achieved sampling rate within 0.01% of nominal, sample loss below 0.02% even at sixteen inlets, and inlet-side pull time indistinguishable between modes. The M4 Pro outlet’s load-regime change between four and eight inlets appears in both modes and is therefore independent of encryption, most plausibly reflecting operating-system network-stack scheduling at higher connection counts.

**Figure 8.**
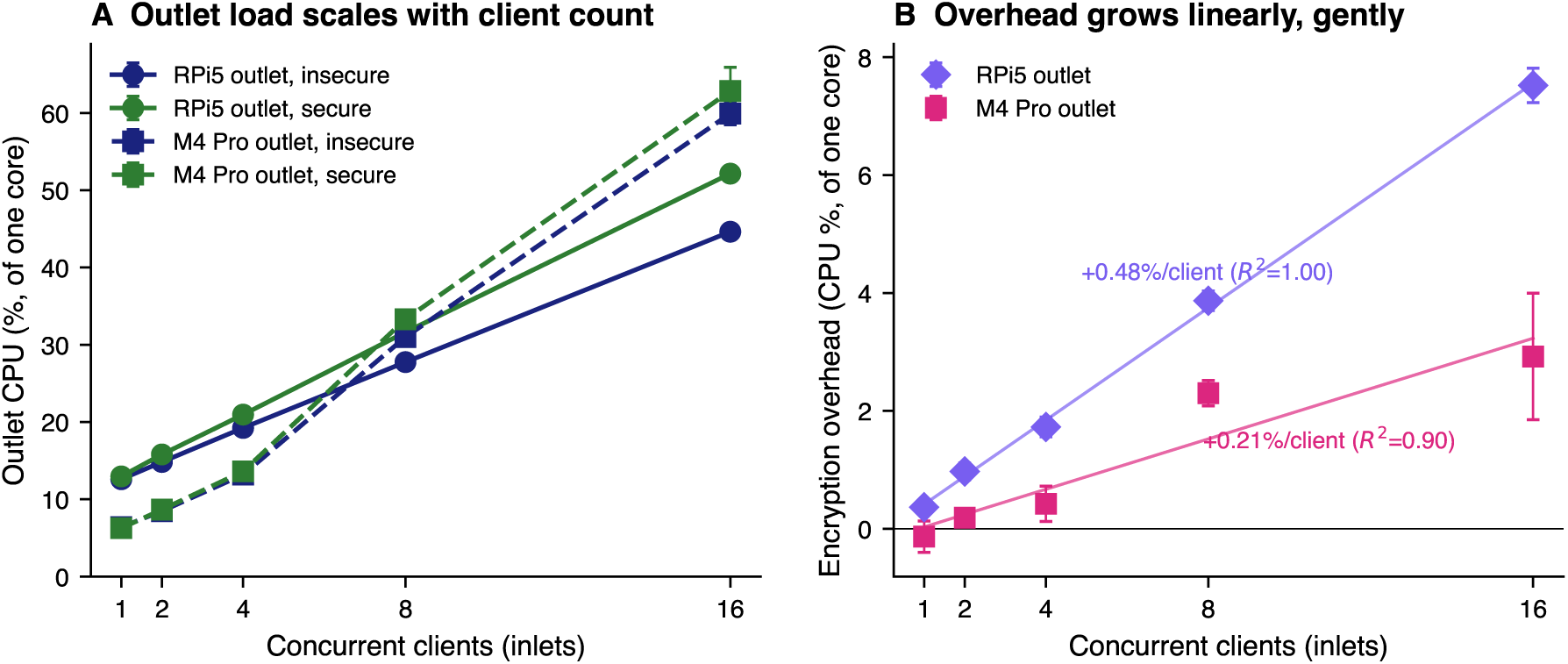
Concurrent-client scalability over a real two-machine Gigabit Ethernet link, with a Raspberry Pi 5 and an Apple M4 Pro Mac Mini each serving as outlet for 1, 2, 4, 8, or 16 inlets at 64 channels and 1000 Hz. CPU is the outlet process’s utilization as a percentage of one core, so the within-machine secure-versus-insecure comparison is the controlled quantity and absolute values are not comparable across architectures. **(A)** Outlet CPU versus client count per machine and mode (error bars: between-block standard deviation); load is dominated by transmission fan-out, and the M4 Pro shows a network-stack regime change near eight clients in both modes. **(B)** Encryption overhead (secure minus insecure outlet CPU) versus client count, with a per-machine least-squares line and slope; the overhead grows linearly at a small per-client rate.

### 4.3 Microcontroller Performance

The ESP32-DevKitC v4 validation characterizes the liblsl-ESP32 reimplementation under the same channel- and rate-sweep configurations as the desktop platforms, scaled to the microcontroller’s resource envelope (Figure 9). Because the ESP32 maintains a monotonic boot-relative clock, we report push-path timing (the cost of an application-level `push_sample()` call on the application core, while encryption and TCP transmission run asynchronously on the second core) and resource utilization rather than end-to-end latency.

**Figure 9.**
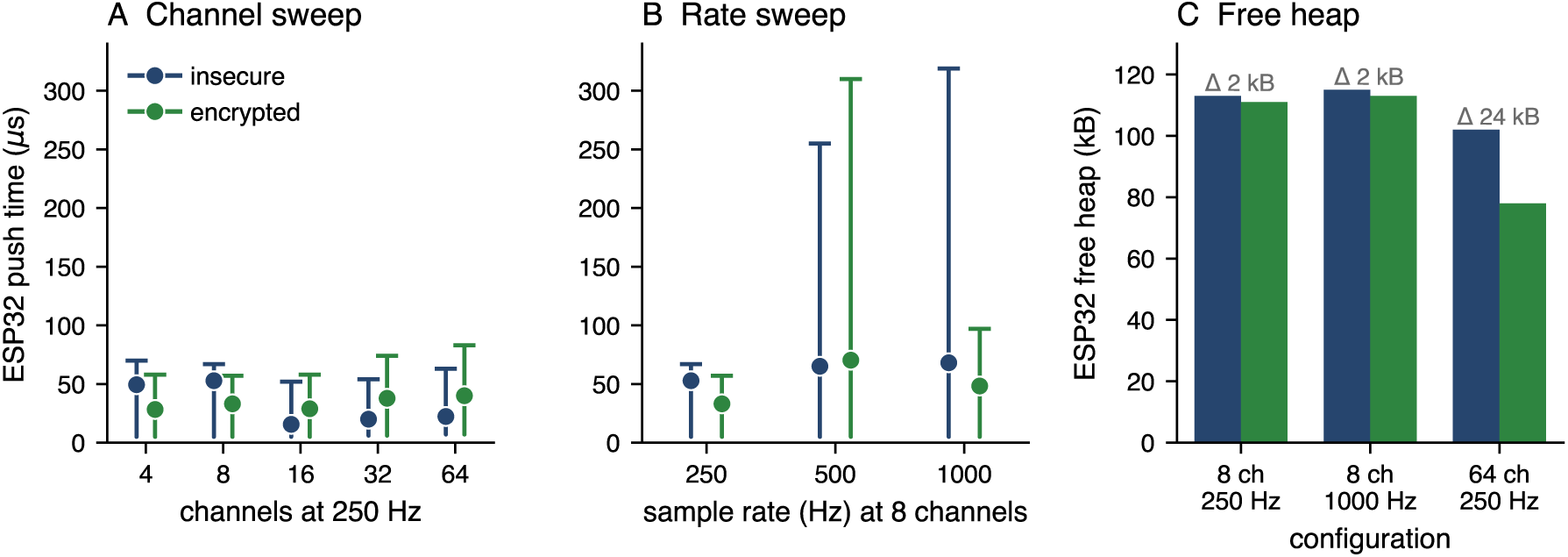
ESP32-DevKitC v4 microcontroller benchmark (Xtensa LX6 dual-core at 240 MHz, 520 kB SRAM, IEEE 802.11n WiFi). **(A)** Push-path timing across a channel sweep at 250 Hz; the marker is the mean push time and the line spans the minimum to the 95th percentile, the heavy WiFi-backpressure tail omitted for legibility. **(B)** Push-path timing across a sampling-rate sweep at 8 channels, same axis. **(C)** Mean free SRAM with and without encryption; the footprint is about 2 kB at 8 channels, growing to about 24 kB at 64 channels as the per-chunk buffers scale with payload size. Running ChaCha20-Poly1305 asynchronously on core 1 renders push-path encryption overhead invisible to the application core in all configurations.

Across the channel sweep at 250 Hz (4–64 channels, float32), mean push times were 16–53 *µ*s insecure and 28–40 *µ*s encrypted, the encrypted mode showing lower variance because the dedicated feed task absorbs WiFi backpressure spikes that otherwise leak to the application path. Across the rate sweep at 8 channels, both modes sustained zero packet loss at 250 and 500 Hz, and at 1000 Hz the encrypted mode showed lower 95th-percentile push times than insecure (97 *µ*s versus 319 *µ*s) and lower packet loss (0% versus 0.02%) because the asynchronous design prevents queue head-of-line blocking under WiFi jitter, consistent with the ring-buffer absorption hypothesis. Encryption increased SRAM consumption by approximately 2 kB at 8 channels (the encrypted-session and authorization-state buffers), growing to approximately 24 kB at 64 channels as the per-chunk buffers scale with payload size. Even in the most demanding 64-channel encrypted configuration the device retained ample free heap (mean 78 kB, transient minimum near 16 kB during connection setup) and sustained the full sweep without packet loss.

Bidirectional encrypted interoperability between the ESP32 firmware and the desktop secureLSL build was verified by streaming identical 8-channel float32 sequences in both directions over the same WiFi network; received samples matched transmitted samples bit-exactly across 30-second windows in both streaming directions.

### 4.4 Interoperability Validation

All four combinations of outlet and inlet languages (Python and MATLAB each paired with both) successfully established encrypted connections and transmitted data with full fidelity, with relative errors below 10^−6^ for floating-point data, confirming that cryptographic operations introduce no precision loss. Coverage included all supported data types (float32, float64, int16, int32, string) and sampling rates from 10 Hz to 10000 Hz. Equivalent automated coverage of the C++ binding is planned for a forthcoming release.

### 4.5 Deployment Simplicity

The change presented to an operator who already runs LSL is deliberately small: source instruments and recording applications require zero code changes, and security is enabled through a single configuration-file addition and a library replacement. Figure 10 summarizes the three artefacts an operator touches: the configuration block that distinguishes a secured install (a), the command sequence that provisions it (b), and the connection-outcome logic that follows (c).

**Figure 10.**
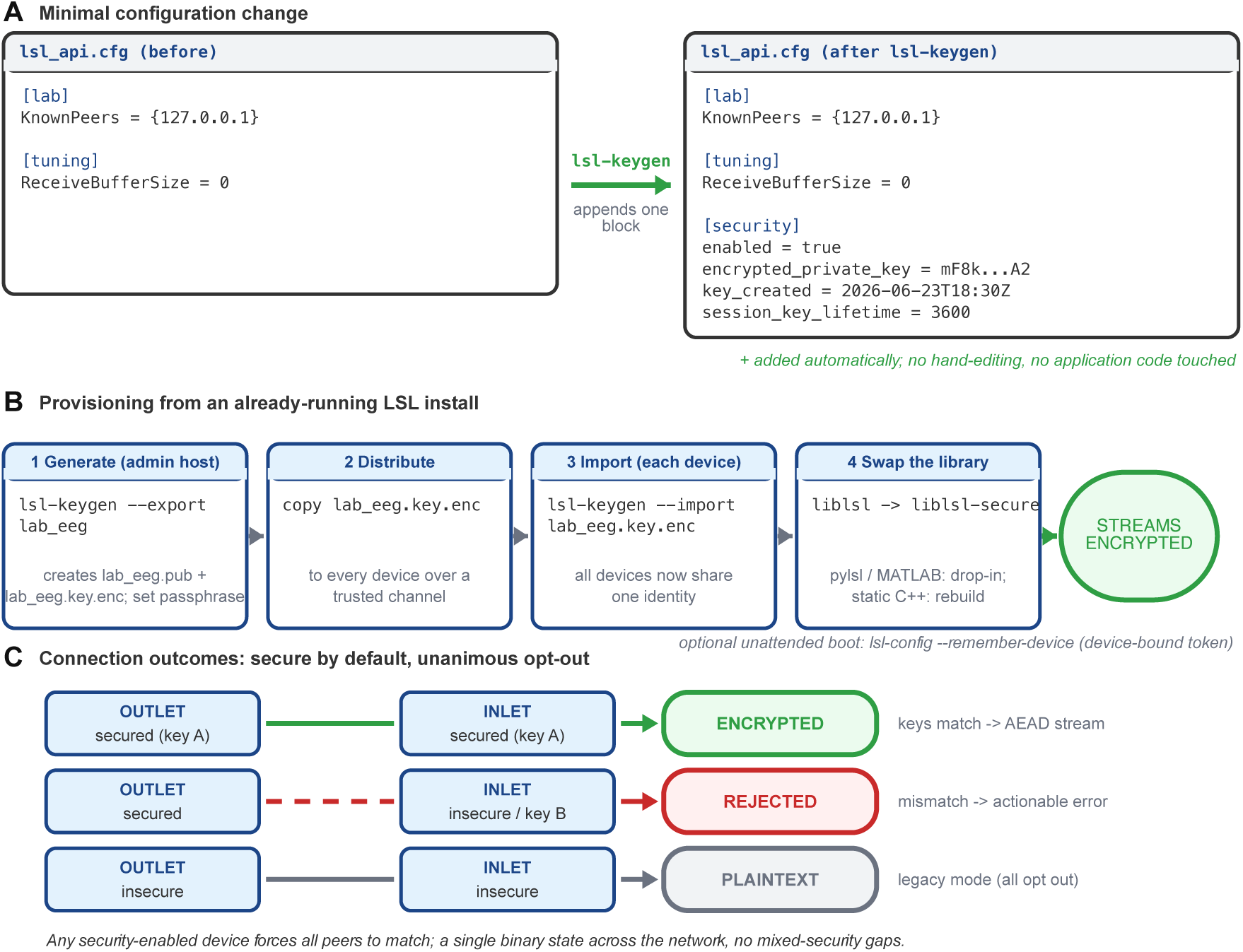
Deploying Secure LSL from an existing installation. **(A)** The only configuration change: `lsl-keygen` appends one `[security]` block to the existing `lsl_api.cfg`, leaving all prior settings untouched. **(B)** Provisioning: an administrator generates and exports a keypair once, the encrypted key file is distributed and imported on every device, and the *liblsl* binary is replaced with *liblsl-secure*, a drop-in change for dynamically linked clients (*pylsl*, MATLAB) and a one-time rebuild for static C++; an optional device-bound token enables unattended boot. **(C)** Connection logic under the secure-by-default, unanimous-opt-out model: matched secured peers stream encrypted data, any mismatch is rejected with an actionable error, and a fully opted-out network falls back to plaintext.

A laboratory reaches encrypted operation in three commands and one library swap (Figure 10b). A trusted administrator runs `lsl-keygen --export lab eeg` once, generating a fresh Ed25519 keypair, prompting for a passphrase, and writing a public file `lab eeg.pub` and an encrypted key file `lab eeg.key.enc`; this file is copied to every device over a trusted channel and imported with `lsl-keygen --import lab eeg.key.enc`, after which all devices share a single identity. Finally the *liblsl* binary is replaced with *liblsl-secure*: dynamically linked clients including the Python (*pylsl*) and MATLAB bindings pick up the change with no recompilation, while statically linked C++ applications are rebuilt once. Unattended sensors can store a device-bound unlock token with `lsl-config` --remember-device instead of entering the passphrase on every start.

The only configuration change is a single [security] block that lsl-keygen appends to the existing lsl api.cfg file (Figure 10a); no existing keys are edited and the remainder of the file is left untouched:

~~~
[security]
enabled = true
encrypted_private_key = mF8k…A2
key_created = 2026-06-23T18:30:00Z
session_key_lifetime = 3600
~~~

The encrypted_private_key field holds the Argon2id-encrypted private key, so a stolen configuration file does not expose the credential; an operator who opts out of passphrase protection instead sees a plaintext private_key field. Because every secured device carries the same block, the connection outcome reduces to a binary decision (Figure 10c): two secured peers whose keys match establish an encrypted stream, any secure/insecure or key mismatch is rejected with an actionable error, and a fully opted-out network continues in plaintext for backward compatibility.

The complete deployment timeline for a typical laboratory is approximately two weeks: one week for key generation and distribution, one day for simultaneous library updates, and one week validating encrypted operation, substantially faster than architectures requiring application-level modifications.

## 5 Discussion

### 5.1 Architectural Trade-offs

The unified model deliberately trades configuration flexibility for deployment simplicity and robustness. Requiring all devices to agree on security mode eliminates partial-security vulnerabilities, prevents downgrade attacks that force insecure connections, and simplifies compliance validation by ensuring a single security posture; binary failure modes eliminate the troubleshooting complexity inherent in systems with multiple security levels or optional encryption.

The primary limitation is coordinated deployment: all devices must update together rather than migrating incrementally. This is arguably a feature, since prolonged mixed-environment transitions create exactly the partial-security vulnerabilities the architecture prevents, and the two-week timeline demonstrated in testing confirms that coordinated migration is feasible for typical research environments.

Metadata exposure is an accepted trade-off: stream names, channel counts, and sampling rates remain visible to network observers even when data is encrypted, analogous to broadcast WiFi SSIDs, so an observer learns that a “64-channel EEG stream at 1000 Hz” exists without accessing brain data. Encrypting metadata would break LSL’s service discovery and require extensive protocol redesign for marginal benefit.

### 5.2 Comparison with Alternative Approaches

We evaluated several alternatives during design. Application-level encryption, with each of the 150+ LSL applications implementing its own security layer, would require years of development across multiple languages with no guarantee of interoperability. Transport-layer security using Transport Layer Security or its datagram counterpart (TLS/DTLS) [27] is a natural candidate, and TLS 1.3’s pre-shared-key and raw-public-key modes (RFC 8446 §2.2 and RFC 7250 [28]) avoid a certificate authority, so certificate infrastructure is not the fundamental obstacle. The difficulties are structural: LSL discovers peers through UDP multicast and combines UDP and TCP transports that do not map cleanly onto a single TLS/DTLS session, retrofitting TLS would break backward compatibility and complicate connection-state management, and the same stack cannot be carried unchanged onto the constrained Xtensa target. Network-layer VPNs such as WireGuard [29] or IPsec/IKEv2 *do* authenticate peers, contrary to a common assumption, and WireGuard is a deliberately simple pre-shared-key design; but a VPN secures the link rather than the stream, requires network-infrastructure changes often beyond laboratory control, leaves the security boundary invisible to the LSL protocol, and does not extend to microcontroller endpoints joining the LSL network directly. Implementing security inside *liblsl* keeps the trust boundary at the stream, makes the secure-versus-insecure state visible and enforceable within the protocol, and ports to the same clean-room microcontroller reimplementation as the desktop hosts.

Optional or opportunistic security that falls back to plaintext would recreate the mixed-security and downgrade vulnerabilities our unified model prevents, and per-stream policies would require extensive application modifications, enable cross-stream attack vectors, and complicate compliance validation. The core-library approach instead keeps security logic in a single auditable codebase rather than scattered across hundreds of applications, so the entire ecosystem gains protection through library updates alone.

### 5.3 Adoption Pathways

Adoption pathways depend on context. New installations enable security from the outset by deploying security-enabled liblsl and generating keys during setup, achieving immediate encrypted operation. Existing deployments migrate through coordinated key distribution and simultaneous library updates on the validated two-week timeline, which regulatory urgency can compress through parallel provisioning. Although each deployment requires coordinated migration, the model’s migration-pressure property, where adding one security-enabled device encourages network-wide adoption, supports gradual organizational rollout, and research groups can pilot security in isolated test environments before committing to production.

### 5.4 Ecosystem Integration

The core implementation resides in a dedicated `secureLSL` repository under the SCCN organization, containing the modified liblsl source, key generation tools, and documentation. Rather than forking ecosystem applications, we contribute security support through pull requests: the pylsl bindings gain automatic preference for `liblsl-secure` when available plus Python wrappers for the version and stream-security APIs, and recording and visualization tools such as LabRecorder and SigVisualizer gain security-status indicators (lock icons for encrypted streams, version information in About dialogs). This integrates security naturally without fragmenting the community across parallel forks. The distinct binary naming (`liblsl-secure` rather than `liblsl`) is both a safety mechanism, preventing accidental linking against the wrong library, and a migration aid, since both versions can coexist during transition.

### 5.5 Licensing and Sustainability

Westner et al. [30], in *Cycling on the Freeway: The perilous state of open-source neuroscience software*, document the systemic underfunding and maintainer attrition that threaten the shared infrastructure on which much of neuroscience research now depends. Permissive open-source licensing has been foundational to the LSL ecosystem and to reproducible neuroscience, but it provides no built-in pathway for recovering the engineering cost of long-term maintenance, security hardening, regulatory tracking, and binding maintenance across languages and platforms. The recent shift toward clinical and commercial deployment of LSL pipelines, together with the regulatory obligations of Section 5.6, sharpens this gap: research-grade infrastructure is increasingly asked to carry production-grade guarantees without a production-grade funding base.

Secure LSL is, to our knowledge, the first deliberate attempt within the neuroscience-software community to address this gap with a *source-available* model, keeping the software fully accessible and free for non-profit academic research, teaching, and training while requiring a commercial license for for-profit and revenue-generating use. The full source, build system, key-generation utilities, benchmark scripts, and figure pipeline are visible and auditable in the public repository, and the license grants non-commercial academic users the right to reproduce and benchmark the results reported here and to modify the code to verify or extend them, so reproducibility, cryptographic peer review, and academic extension are not gated; only commercial use and competitive benchmarking require a separate license. The commercial path captures value from industrial users who derive direct economic benefit and cycles it back into maintenance, security review, post-quantum migration, regulatory tracking, and ecosystem integration. We view this as a first experiment rather than a fixed answer, hoping that source-available and dual licensing become a normal part of the neuroscience software toolbox alongside the permissive licensing that built the field.

### 5.6 Regulatory Landscape and Compliance

Section 1.2 introduced the regulations that make plaintext biosignal transport untenable for clinical and multi-institution deployments; we now examine the specific obligations and how the architecture addresses them. The discussion concerns identifiable biosignal data linkable to individuals; de-identified data (e.g., under HIPAA Safe Harbor at §164.514(b)) and properly anonymised data (under GDPR Recital 26) fall outside these regulations, though encryption remains best practice against re-identification and as protection for proprietary paradigms.

Two framing points apply throughout. First, the architecture supplies cryptographic *mechanisms* that support these obligations but does not by itself confer legal compliance, which also depends on organizational policy, physical security, access governance, and data-handling practices outside the transport layer. Second, the default configuration prioritizes zero-configuration usability and is explicitly not a turn-key compliance profile: regulated and clinical deployments are expected to enable the stricter configuration (passphrase-protected keys, which add the multi-factor knowledge factor; device-bound session tokens; security-event logging; and network-wide secure mode) and pair it with the organizational controls each framework requires. The mapping below notes where the supporting configuration is non-default.

#### 5.6.1 United States

##### HIPAA

The HIPAA Security Rule designates encryption of electronic Protected Health Information (ePHI) in transit as an addressable specification under 45 CFR §164.312(e)(2)(ii). Though addressable rather than mandatory, encryption is expected by regulators for any networked ePHI transmission, with Security Rule penalties reaching over US$2 million annually for willful neglect [3]. Research-only devices are generally exempt from FDA medical-device clearance, but FDA cybersecurity guidance applies once the equipment is used clinically. The architecture supports the HIPAA Technical Safeguards through access control restricting the secured network to credential holders (§164.312(a)(1)), audit controls via security event logging (§164.312(b)), person-or-entity authentication via verification of the shared credential (§164.312(d)), and transmission security via ChaCha20-Poly1305, an Internet Engineering Task Force (IETF) standardized algorithm [21] (§164.312(e)). Where the unique user-identification element is required, that identity is supplied by the host application and organizational directory rather than the shared transport credential, which certifies group membership rather than individual device identity.

#### 5.6.2 European Union

##### NIS2, EHDS, GDPR, and CRA

The Network and Information Security Directive 2 (NIS2), effective October 18, 2024, establishes cybersecurity risk management measures and reporting requirements for healthcare and research institutions [5]. NIS2 mandates multi-factor authentication and encryption where appropriate, with sanctions reaching €10 million or 2% of worldwide turnover for non-compliance [6]. The European Health Data Space (EHDS) regulation, in force since March 26, 2025, targets health-data exchange and requires organizations to guarantee the security, integrity, and confidentiality of health data, including encryption of data in transit and strict access controls [7]. GDPR Article 32 continues to require “appropriate technical measures” including encryption of personal data, with fines reaching €20 million or 4% of global annual revenue [4]. The Cyber Resilience Act (CRA), in force since December 10, 2024, requires that products with digital elements protect the confidentiality and integrity of data, including data in transit, with full compliance required by December 2027 [8] and is particularly relevant for commercial neurotechnology products and research instruments that incorporate networked data streaming. Biosignal data carrying participant identifiers, particularly EEG and electromyography (EMG), constitutes personal data under all four frameworks.

Across all four EU frameworks the architecture provides the same core mechanisms: ChaCha20-Poly1305 encryption of health and personal data in transit, access control via authenticated device connections, Poly1305 integrity guarantees, and security event logging supporting the mandated reporting timelines (Table 3), with the GDPR Article 32 obligation to test security-measure effectiveness met by the validation procedures of Section 3.10 and the CRA’s 24-hour notification supported by tamper-detection and connection-rejection logging. The NIS2 multi-factor-authentication expectation is satisfied only when passphrase-protected keys are enabled, a non-default configuration.

**Table 3:** Global regulatory-compliance mapping for the Secure LSL architecture.

| Regulation | Jurisdiction | Encryption | Authentication | Audit logging | Breach reporting |
| --- | --- | --- | --- | --- | --- |
| HIPAA | United States | §164.312(e) | §164.312(d) | §164.312(b) | Required |
| NIS2 | European Union | Required | MFA mandated | Required | 24–72 h |
| EHDS | European Union | Required | Required | Required | Per GDPR |
| GDPR | European Union | Art. 32 | Art. 32 | Art. 5(2), 32 | 72 h |
| CRA | European Union | Art. 6 | Art. 6 | Art. 6 | 24 h |
| PIPL | China | Required | Required | Required | “immediately” |
| APPI | Japan | Required* | Required | Required | reduced if encrypted |
| PIPA | South Korea | Required | Required | Required | 72 h |

\*Encryption to high standards reduces breach-reporting obligations under APPI.
*Note:* entries indicate which cryptographic mechanism supports a given obligation; they are not legal determinations. The architecture supplies technical measures that support compliance but does not by itself confer legal compliance, which also depends on organizational, physical, and data-governance controls outside the transport layer, and was not reviewed by legal counsel.

#### 5.6.3 China

##### PIPL, CSL, and DSL

China’s three-pillar framework comprises the Personal Information Protection Law (PIPL), the Cybersecurity Law (CSL), and the Data Security Law (DSL), all effective since 2021 [9]. PIPL classifies biometric characteristics as Sensitive Personal Information requiring enhanced technical protection such as encryption and de-identification, with penalties reaching ¥50 million (approximately US$7 million) or 5% of a company’s China revenue; its stringent data-localisation requirements have led the German Research Foundation, Swedish Research Council, and Swiss National Science Foundation to avoid co-funding projects in China since 2021 [10]. The architecture’s encryption and authentication constitute the mandated technical measures for PIPL Sensitive Personal Information and address CSL network-security requirements against data leakage and theft.

#### 5.6.4 Japan

##### APPI

Japan’s Act on the Protection of Personal Information (APPI), amended in 2020 and 2024, classifies medical and health information as “special care-required personal information” [11]. APPI mandates encryption of sensitive data in transit and at rest to strong standards such as AES-256, and reduces breach-reporting obligations when data is encrypted to the highest standards. The architecture’s ChaCha20 cipher provides 256-bit keys with security equivalent to AES-256, which may support qualification of the secure-by-default mode for APPI’s reduced reporting regime, subject to confirmation under current APPI guidance.

#### 5.6.5 South Korea

##### PIPA

South Korea’s Personal Information Protection Act (PIPA), amended in March 2024, governs personal data across all sectors and extends to offshore processing of Korean individuals’ data [12]. Healthcare and medical data are sensitive information, triggering mandatory 72-hour breach reporting to the Personal Information Protection Commission, and the Bioethics and Safety Act requires Institutional Review Board approval for human-subjects research, including pseudonymised data. The architecture addresses PIPA through encryption, authentication preventing unauthorized access, and security event logging enabling the 72-hour reporting, and supports Bioethics and Safety Act requirements for both identified and pseudonymised research data.

### 5.7 Limitations and Future Work

Several limitations suggest future directions. The current per-device key generation and distribution, while straightforward for typical laboratories, becomes administratively burdensome beyond 100 devices, where automated provisioning integrating enterprise key management platforms such as HashiCorp Vault or cloud services would help. Post-quantum cryptography is a longer-term consideration: while Ed25519 and ChaCha20-Poly1305 are strong against classical computers, a sufficiently large quantum computer could break the Ed25519 signatures. NIST finalized the first post-quantum standards in 2024, including ML-KEM (a module-lattice key-encapsulation mechanism) and ML-DSA (a module-lattice digital signature algorithm) [31]; once efficient implementations are widely available, a migration path should be developed, which the modular cryptographic architecture facilitates by isolating algorithm-specific code. Encrypted storage also remains out of scope: the architecture secures data in transit but not at rest, so Extensible Data Format (XDF) files remain unencrypted unless applications add their own file encryption, which future work could integrate through filesystem-level or application-level encrypted containers. Finally, the security validation reported here establishes implementation correctness and conformance to the intended rejection behavior, but the custom signed-ephemeral-X25519 handshake has not yet undergone independent red-team assessment, protocol fuzzing, timing side-channel measurement of the compiled binaries, or formal or symbolic protocol verification of the kind applied to protocols such as WireGuard; we regard these as important next steps before unsupervised clinical deployment.

### 5.8 Impact on LSL Ecosystem

This work expands LSL beyond research environments into domains requiring robust data protection. Clinical brain-computer interfaces and neurofeedback systems can now use LSL while meeting requirements across the major jurisdictions (Table 3), and multi-institution collaborations can deploy shared infrastructure without exposing proprietary data to partners, enabling federated analysis previously impractical, especially given that funding agencies in Germany, Sweden, and Switzerland have restricted co-funding over data-protection concerns. Commercial manufacturers serving international markets can deploy a single security implementation rather than separate per-region compliance solutions, and the clean-room `liblsl-ESP32` implementation extends this reach to wearable, ambulatory, and home-monitoring biosensors that cannot host the upstream liblsl C++ codebase, while preserving wire-protocol compatibility and the unanimous-security guarantee.

The architecture preserves LSL’s core strengths (zero-configuration networking, automatic device discovery, seamless reconnection, and millisecond synchronization) while eliminating its critical limitation of plaintext transmission. Implementing security within the protocol core rather than across the ecosystem’s applications enables broad adoption without burdening the diverse community of LSL developers, yielding a platform suitable for research and clinical environments while remaining accessible to the casual user who values LSL’s simplicity.

## 6 Conclusion

We present a unified security architecture for the Lab Streaming Layer *synchronous data collection* framework that achieves encrypted biosignal streaming through transparent core library modifications. It combines a shared Ed25519 keypair for authorization with ChaCha20-Poly1305 authenticated encryption, implemented within liblsl so all LSL applications gain security without code changes. The “secure by default with unanimous opt-out” model enforces network-wide consensus through public key verification, eliminating the vulnerabilities and complexity of mixed-security environments while providing clear failure modes. Argon2id-encrypted key storage and device-bound session tokens raise the bar against unauthorized credential transfer, though they cannot protect a host whose memory is already compromised.

Validation across desktop, laptop, embedded ARM, and Xtensa microcontroller platforms demonstrates sub-millisecond added latency in all desktop and ARM configurations, with single-digit percentage overhead for the standard 64-channel, 1000 Hz configuration (6.4% on the M1, 7.4% on the i9-13900K, and 8.5% on the Raspberry Pi 5, and an upper bound of 4.2% on the M4 Pro, where the difference is not statistically distinguishable from zero at the block level). The *liblsl-ESP32* implementation exhibits no measurable push-path overhead through dual-core asynchronous encryption and adds approximately 2 kB of SRAM, extending operation to wearable and ambulatory biosensors. Packet loss was zero across the benchmark blocks on all desktop and ARM platforms and at 250 and 500 Hz on the ESP32, with full Python/MATLAB interoperability and 100% detection of tampering and replay attacks. The architecture addresses regulatory requirements across HIPAA (United States); NIS2, EHDS, and GDPR (European Union); PIPL (China); APPI (Japan); and PIPA (South Korea), supporting compliant international collaborations on an approximately two-week deployment timeline.

By implementing security within the protocol core and prioritizing simplicity and robustness over configuration flexibility, this work transforms LSL from a research-only protocol into a security-capable platform for clinical neurotechnology, multi-institution collaborations, and commercial products, while preserving the zero-configuration simplicity the neuroscience community relies on.

## Competing Interests

Following institutional review, The Regents of the University of California elected to file a patent application covering the encryption architecture described in this work and to protect the associated software copyright; the patent and license therefore belong to The Regents of the University of California, on which the authors are named inventors. The technology is available under commercial licensing terms for industry use, as described below. Consistent with journal data-and-materials-sharing policy, the license permits non-commercial academic users to reproduce, benchmark, and modify the software to verify or extend the results reported here; only commercial use and competitive benchmarking are reserved to the license holder.

## Use of Agentic Tools

In the interest of transparency, we note that the security architecture and algorithm design reported here are human-conceived and human-owned, while the supporting engineering workflow, including code implementation, testing, and benchmark orchestration, was facilitated by agentic artificial-intelligence coding tools under continuous human direction and review, supported by an open, auditable framework of structured agent workflows [32]. We see such tools accelerating scientific work but believe they warrant stronger checks and balances than are presently standard. The authors assume full responsibility and authorship for this manuscript and the Secure LSL software, including all cryptographic claims, source code, and reported results.

## Acknowledgments

This project was made possible by the Swartz Gift to the University of California, San Diego, and by the Swartz Center for Computational Neuroscience (SCCN), Institute for Neural Computation, UC San Diego, which has maintained the Lab Streaming Layer project and provided technical consultation throughout this work. The cryptographic architecture and protocols described here are the subject of a pending patent application owned by The Regents of the University of California; the software is distributed under a source-available license that grants free non-commercial academic use and reserves commercial use to the copyright holder, with the full `LICENSE` text in the `secureLSL` repository.

## Funding

Swartz Gift to the University of California, San Diego; Swartz Center for Computational Neuroscience (SCCN), Institute for Neural Computation, UC San Diego.

## Author contributions

**Seyed Yahya Shirazi:** Conceptualization, Methodology, Software, Validation, Formal analysis, Investigation, Data curation, Visualization, Writing – original draft, Writing – review & editing, Project administration. **Scott Makeig:** Funding acquisition, Supervision, Visualization, Writing – review & editing.

## Data availability

The Secure LSL reference implementation and documentation are available at https://github.com/sccn/secureLSL and archived on Zenodo [33], including the desktop and embedded ARM `liblsl-secure` build, the `liblsl-ESP32` clean-room microcontroller reimplementation, key generation utilities (`lsl-keygen`, `lsl-config`), and the MkDocs documentation site. The secure library binary is named `liblsl-secure` to prevent confusion with the original `liblsl`, and the microcontroller firmware is namespaced under `liblsl-ESP32` for the same reason. Ecosystem integration is provided through pull requests to existing LSL application repositories (pylsl, LabRecorder, SigVi-sualizer). Performance benchmarks, validation test suites, raw per-platform JSON results, and the figure-generation scripts used to produce all benchmark figures in this manuscript are available in the repository. The software is freely available for nonprofit academic research use, including reproducing and benchmarking the results reported in this manuscript and modifying the code to verify or extend them; commercial use and competitive benchmarking require a separate license. As Westner et al. [30] document, maintaining critical neuroscience research infrastructure faces persistent sustainability challenges; the dual-licensing approach aims to ensure continued development by enabling commercial users to contribute to the ecosystem’s long-term viability while keeping the software accessible to the research community.

